# Rapid evolution in response to climate-change-induced drought and its demographic and genetic controls

**DOI:** 10.1101/2022.05.10.491393

**Authors:** John W. Benning, Alexai Faulkner, David A. Moeller

## Abstract

Populations often vary in their evolutionary responses to a shared environmental perturbation. A key hurdle in building more predictive models of rapid evolution is understanding this variation – why do some populations and traits evolve while others do not? We combined long-term demographic and environmental data, estimates of quantitative genetic variance components, a resurrection experiment, and individual-based evolutionary simulations to gain mechanistic insights into contrasting evolutionary responses to a severe multiyear drought. We examined five traits in two populations of a native California plant, *Clarkia xantiana*, at three timepoints over 7 years. Earlier flowering phenology evolved in only one of the two populations, though both populations experienced similar drought severity and demographic declines and were estimated to have considerable additive genetic variance for flowering phenology. Pairing demographic and experimental data with evolutionary simulations suggested that while seed banks in both populations likely constrained evolutionary responses, a stronger seed bank in the non-evolving population resulted in evolutionary stasis. Gene flow through time via germ banks may be an important, underappreciated control on rapid evolution in response to extreme environmental perturbations.

## Introduction

Extreme environmental perturbations offer excellent opportunities to examine evolution in natural populations on ecological timescales [1]. Rapid evolution in response to strong episodes of environmental change has been documented in many taxa, including lizards [2,3], finches [4], and plants [5]. Beyond offering fundamental insights into the evolutionary process, this and related work speak to the potential for natural populations to evolve in response to anthropogenic forcings like climate change, and to the importance of adaptation for the persistence of populations [6–8]. However, it remains poorly understood as to why rapid evolution often fails to occur in some populations despite the same environmental episode resulting in evolution in other nearby populations.

Resurrection experiments are increasingly used to directly examine evolutionary change before and after an environmental event predicted to exert strong selection on populations [5,9]. The power of resurrection experiments lies in their ability to precisely quantify short-term evolutionary responses, rather than only making predictions of the trajectory of evolution, while controlling for confounding environmental effects on phenotypes. In short, the resurrection approach entails rearing different generations of a population contemporaneously in a common environment to directly quantify phenotypic evolution. Though only a modest number of resurrection studies have been published to date [10], such experiments with plants have sometimes demonstrated rapid evolution in response to periods of drought [5,11,12].

Just as important as these positive results, however, are instances in which resurrection experiments do *not* find evidence of rapid evolution. Populations of a species can vary in their responses to the same climatic fluctuation [12–14], and even extreme climatic anomalies can result in no discernible phenotypic evolution of putatively important traits [15]. What underlies this variation in populations’ evolutionary responses to extreme events? While the resurrection approach is a powerful method to determine if phenotypic evolution has occurred, it does not typically provide insight into the demographic and genetic controls on evolutionary change.

A more comprehensive understanding of the causes and consequences of rapid evolution can be gained by coupling evolutionary analyses with long-term demographic and quantitative genetic investigations. Sharp declines in population size due to environmental perturbations will reduce the efficacy of selection and increase the influence of stochastic processes like genetic drift [16,17]. Such demographic, environmental, or genetic stochasticity may stymie adaptation, potentially causing population extinction before ‘evolutionary rescue’ can occur [18–20]. Models and experiments have further shown that evolutionary rescue is less probable when the environment changes suddenly compared to when the change is gradual [21]. As such, the rate and severity of demographic decline can provide insight into instances where some populations fail to evolve in response to an extreme event that drove rapid evolution in other conspecific populations.

For organisms with dormancy, long-term outcomes of selection can also be affected by the presence of a germ bank (seed or egg bank). Populations that often experience significant environmental fluctuations may evolve mechanisms that confer dormancy as a bet-hedging strategy [22]. For annual plants, seed banks thus result in overlapping, instead of discrete generations. While germ banks may buffer populations from extinction [23,24], they may also constrain evolutionary responses to environmental change due to gene flow among generations [25,26]. This “temporal migration” can slow the rate of adaptive evolution [27,28]. Alleles favored prior to a selective event will be ‘reintroduced’ to the population from the germ bank during and after the event. If those same alleles are selected against during the event, this gene flow can retard adaptation [but see 29]. Moreover, the environmental and demographic history of populations just prior to an extreme event likely affects the magnitude of gene flow from the past. For example, high fecundity of generations immediately prior to an extreme environmental event would increase input to the seed bank. Therefore, historical demographic data can also provide insight into the influence of germ banks on population responses to selection.

Strong environmental perturbations typically exert selection on traits, but the same perturbation may still result in selection regimes that vary among sites [5]. The response to any selection pressure will depend upon the presence of sufficient additive genetic variance for traits underlying fitness variation [30,31]. A population’s response to selection will be directly proportional to a given trait’s narrow-sense heritability, *h*^2^ = *V_A_/V_p_* [32]. In the context of resurrection experiments, measuring the additive genetic variance of ecologically-important traits may help us understand why traits do or do not evolve in different populations exposed to similar selective regimes. While tests of rapid evolution are increasingly common, few studies have simultaneously examined quantitative genetic variation and its potential role in facilitating or hindering adaptation [but see 5].

Furthermore, resurrection experiments usually examine cohorts from before and at the end of an extreme event [but see 33]. A key question is how long evolutionary changes that occur in response to these events persist beyond them. Including later cohorts in a resurrection experiment allows us to assess the temporal stability of evolutionary responses and the extent to which dormancy might delay responses to selection. When paired with associated demographic data from natural populations, such a design gives inference to how rapid evolution (or its absence) influences a populations’ longer-term phenotypic and demographic trajectory.

The Southwest of the United States is in the midst of the most severe multi-decadal drought (i.e., megadrought) in recorded history (at least since 800 C.E.) [34,35]. Recent analyses have shown that anthropogenic climate change has been a key driver of this drought and accounts for an estimated 42% of the soil moisture anomaly [35,36]. We examined evolutionary responses to the most severe episode of this climate anomaly, which began in the latter half of 2011 and ended in late 2015 in our study area. Our work focused on a well-studied annual plant, *Clarkia xantiana* ssp. *xantiana*, which is endemic to Southern California, U.S.A. We used a resurrection experiment to test whether traits likely to mediate drought adaptation exhibited rapid evolution and whether genetic or demographic constraints modulated responses.

We monitored environmental and demographic variation over 12 years (2006 – 2017), spanning a period from six years prior to drought until two years after the drought ended. Those data provided insight into the severity and rate of environmental and demographic change, as well as the magnitude of input to the seed bank in years just prior to the drought episode. To test for rapid evolution in response to drought, we used a resurrection experiment with pedigreed individuals from three time points — prior to the prolonged drought (2011), at the end of the drought (2015), and two years later, after more average precipitation resumed (2017). This latter sample allowed us to examine whether any evolutionary responses persisted, increased, or evolved back toward the original population mean. We measured a suite of traits that prior studies have shown to confer drought escape and/or avoidance, and estimated additive genetic variance for each of these traits in each population. Lastly, we leveraged these demographic and experimental data in the construction and analysis of an individual-based evolutionary simulation model to test the hypothesis that a seed bank influenced the evolutionary responses of these populations to extreme drought.

## Material and Methods

### Study System

*Clarkia xantiana* ssp. *xantiana* A. Gray (Onagraceae) is a predominantly outcrossing winter annual native to the southern Sierra Nevada foothills and Transverse Ranges of California, USA [37]. In this Mediterranean climate, the bulk of precipitation occurs during winter and early spring. Plants germinate November - March during the rainy season, begin flowering in May, and set seed in late June. In this study, we focus on two populations, KYE and S22. KYE occupies oak woodland habitat on granite-derived soils characteristic of the more mesic, western portion of *C. x. xantiana’s* distribution (35.6240674°, −118.5156798°). S22 is located near the subspecies’ eastern geographic range limit and occupies a more xeric, higher elevation site in pine woodland on metasedimentary-derived soils (35.83996°, −118.450386°). These metasedimentary-derived soils occur along the Kern Canyon Fault, a ca. 150km fault that parallels the Kern River through the Southern Sierra [38,39]. [Site identifiers are 57x (KYE) and 22x (S22) for consistency with previous and future *C. x. xantiana* studies).] Though it was long assumed that *Clarkia* species had little if any seed storage [e.g., 40], recent work has shown that *C. x. xantiana* populations can harbor substantial seed banks [41,42].

### Climatic and demographic data

We obtained growing season precipitation data (November - June) for years 2006-2017 from long-term weather monitoring stations (HOBO Onset) at each site. Demographic data were collected as part of a long-term study of 36 populations across the range of *C. x. xantiana* in years 2006-2017 (detailed in Supp. Mat. A). Using a population projection model approach, we combined estimates of seed input in each year with estimates of seed bank vital rates (survival and germination) for KYE and S22 from [42] to estimate the age distribution of germinants in each population in each year (assuming a maximum seed age of 3 years). Siegmund et al. (2022) estimated these vital rates for the two populations in our study as: seed survival rate (KYE = 0.66; S22 = 0.61), germination rate (KYE = 0.13; S22 = 0.27). The multi-year field experiments used to estimate these vital rates are detailed in [42] and [41].

### Seed collection and refresher generation

We used seeds collected as part of the aforementioned long-term demographic study on *C. x. xantiana*. Seeds were collected from a haphazard sampling of plants (dozens to hundreds of plants, depending on field conditions) across the spatial extent of sites KYE and S22 in late June of years 2011 (pre-drought), 2015 (end of drought), and 2017 (after two years of more average precipitation following the drought). All seeds from a given dam constitute a maternal family. Because *C. x. xantiana* is a highly outcrossing species [43] with multiple paternity (DM, unpubl. data), each maternal family sampled from the field was a collection of full and half-sibs. Seeds were stored at room temperature in plastic boxes containing silica desiccant, with maternal families each in their own coin envelope. To standardize maternal environmental effects, we grew plants from each year together in a greenhouse for one “refresher” generation to produce seeds for the resurrection experiment [10].

For each of the three generations of each population (six “cohorts”), we grew 1-5 plants from each of 20 haphazardly selected maternal families (N = 219 plants total) in a fully randomized design in the greenhouse in spring 2018. Most (66%) of maternal families were represented by two plants; representation of families varied due to unequal germination. Within each cohort, half the plants were randomly assigned as sires and each sire was mated to two unique dams (with all plants serving as dams) to produce a pedigreed population for the measurement round of the experiment. We also grew individuals from a third site, Mill Creek, during this generation prior to the resurrection experiment, but dropped this site from the subsequent experiment for feasibility; data from the Mill Creek refresher generation did not suggest evolution of any of the three measured traits (growth rate, days to flowering, SLA; Fig. S4).

### Measurement generation

With the seeds produced from crosses, we grew the six offspring cohorts in a fully randomized design in the greenhouse in spring 2019 to assess phenotypic changes across generations for each population. Of the original 20 maternal families planted for each year cohort in the refresher generation, 16-20 were represented (as sire and/or dam) in this pedigreed population of offspring. We sowed six seeds for each of 22-28 dams from each refresher cohort in plug trays with germination mix in the growth chamber and scored germination for 33 days. There were 83-122 germinants per cohort (Table S2); germination rates ranged from 0.61 to 0.75 (Table S1). Up to four seedlings per dam (haphazardly chosen) were transplanted into individual 656 mL Deepots (Stuewe & Sons, Tangent, OR) filled with a 1:1 mix of sand and potting soil. Pots were arranged in a completely randomized design in the greenhouse on a 16/8 hr light schedule and watered as needed. Final sample sizes varied due to unequal seed availability and germination among cohorts (*n* = 65 - 97 individuals per year cohort for post-germination traits; Table S2).

### Trait measurements

We measured five traits that have previously been shown to be related to drought escape and/or avoidance: days to germination, growth rate, days to flowering, specific leaf area (SLA), and leaf succulence. Faster germination, growth, or flowering phenology may allow plants to take advantage of the relatively mesic early growing season and complete their life cycle before the onset of drier conditions [5,11,44], though delayed germination may also be selected for in arid environments [e.g., 45]. Lower SLA and increased leaf succulence can increase water use efficiency, and evolution in these physiological traits may represent drought avoidance adaptations [44,46]. We also measured two fitness proxies: total number of flowers produced and shoot (aboveground) biomass.

Days to germination was the number of days elapsed between sowing and cotyledon emergence. Growth rate was measured as the number of leaves produced per day, measured for plants 41-43 days post-germination (i.e., growth rate was calculated as the number of leaves at measurement divided by the plant age in days at measurement). One fully-expanded leaf was collected from each plant 64-75 days post-germination; we recorded fresh and dry weight for that leaf and used ImageJ [47] to measure leaf area from a photo of the fresh leaf. SLA was calculated as mm^2^ mg^-1^; leaf succulence was calculated as leaf wet weight / leaf dry weight. Days to flowering was measured as the days elapsed between seed sowing and the opening of a plant’s first flower. Measuring flowering time in such a way, as opposed to the days elapsed between germination and flowering, incorporates variability in time to germination and is more directly relevant to phenological timing as measured in wild populations; both traits showed a similar pattern across years (Table S1). Total number of flowers produced was measured when all flowers had opened on a plant. Shoot biomass was collected at the end of the experiment and plants were dried before weighing. Two pairs of traits were strongly correlated [number of flowers produced and shoot biomass (r = 0.72); SLA and leaf succulence (r = 0.81); Fig. S5]. We report on number of flowers produced and specific leaf area below, as flowers produced is a more direct proxy for plant fitness than shoot biomass, and SLA is a more commonly-reported trait than leaf succulence. We include descriptive statistics for shoot biomass and leaf succulence in Table S1. Our analyses of invisible fraction bias [sensu 48] indicated that any bias in estimation of trait means due to unequal seed survival among genotypes was likely minimal (Supp. Mat. B).

### Statistical analyses for resurrection experiment

All statistical analyses were conducted in R version 4.1.2 [49]. Data were organized and summarized using the tidyr [50] and dplyr [51] packages and plotted using ggplot2 [52]. All data and code necessary to reproduce these analyses are uploaded with the submission and will be archived at FigShare upon acceptance.

#### Phenotypic evolution

We used linear mixed models [package lme4 [53]] to test for differences among year cohorts in the measured phenotypic traits (days to germination, growth rate, flowering date, SLA, and flower number). We analyzed populations separately because our main interest was in differences among year cohorts, not populations, and we had limited power to detect interactions between population and year. All models took the general form of *response ~year + transplant age + (1/sire/dam*). Transplant age (days elapsed between germination and transplanting from germination tray to full pot) was included as a covariate to account for differing seedling ages when transplanted to pots. Sire, and dam nested within sire, were included as random effects to account for non-independence amongst related individuals. For days to germination, dam average seed weight was included as a covariate and transplant age was not. Only seeds that germinated were included in days to germination analyses to avoid any confounding of dormancy with seed viability. Days to germination and SLA were log transformed prior to analysis to better meet assumptions of homoscedasticity and normality of model residuals. We tested whether inclusion of terms improved model fit with Wald λ^2^ tests using Type II SS [Anova.mermod() function in the car package [54]]. If year was a significant predictor at *α* = 0.05, we followed up with pairwise Tukey contrasts between year cohorts using the emmeans package [55]. We computed estimated marginal means for each year cohort using emmeans.

#### Genetic variance components

We estimated quantitative genetic variance components for all traits in both populations using an animal model implemented in the MCMCglmm package [56]. In short, the animal model approach comprises a linear mixed effects model with an individual’s breeding (i.e., additive genetic) value modeled as a random effect [57,58]. A pedigree of the population provides an expectation of how breeding values should covary between individuals, and thus allows for an estimate of a trait’s additive genetic variance in that population. MCMCglmm uses a Markov Chain Monte Carlo approach in a Bayesian framework to approximate the posterior distribution of quantitative genetic variance components (analyses detailed in Supp. mat. C). Following [59] we obtained estimates of additive genetic variance (V_A_), residual variance (V_R_), and the variance explained by fixed effects (V_F_) from the model, and calculated *h^2^* = V_A_ / (V_A_ + V_R_ + V_F_); we report the mean of the posterior distribution of *h*^2^ as our estimate. We considered a trait to exhibit significant heritability if its 95% credible interval for *h*^2^ did not touch 0.001 (in MCMCglmm, variance estimates must be greater than zero). MCMCglmm diagnostic plots and posterior distribution of model parameters are included in SI: MCMCglmm.

### Simulation model

We used an individual-based, genomically-explicit evolutionary simulation model, paired with Approximate Bayesian Computation, to test the hypothesis that seed bank dynamics influenced evolutionary patterns in our populations. We focused this investigation on one trait, flowering phenology, which displayed contrasting evolutionary responses in our two populations (Results). Using the SLiM modeling framework [60], we built a model tracking the demography and evolution of a population of diploid, hermaphroditic individuals that experience an environmental perturbation. Individual female fitness (here, the probability of surviving to set seed) is determined by the deviation of an individual’s phenotype (here, flowering phenology, which is under stabilizing selection and controlled by multiple QTL) from the optimal phenotype; the optimal phenotype is assumed to change based on current environmental conditions. We focus on female fitness (i.e., siring success is not affected by an individual’s phenotype) because earlier work in *C. x. xantiana* has demonstrated strong selection on flowering phenology via female fitness in arid conditions [61] but we have no such tests of selection via male fitness. After a long burn-in period with a relatively stable environment (and thus phenotypic optimum), the model environment and population size are explicitly tied to our observed 12 year data set. There is an environmental perturbation (e.g., drought) in year 7 that changes the phenotypic optimum, and modeled population sizes in each year are proportional to the observed census sizes of natural populations in each year. Full model details are in Supp. Mat. D.

We used Approximate Bayesian Computation (ABC) to 1) test the hypothesis that seed banks influenced evolutionary trajectories in our populations, 2) test the hypothesis that the phenotypic optimum returned to its pre-drought state after drought, and 3) estimate seed bank demographic (seed survival and germination) and evolutionary (change in phenotypic optimum during drought) parameters in both populations. ABC is a statistical method used to estimate model parameters and choose among a set of candidate models when likelihood calculations are intractable [62]. First, many simulations are run where values of the parameters of interest are randomly drawn from their defined prior distributions (Table 1). Then, summary statistics are calculated for each simulation run and are used to compare simulation outcomes with observed data, in order to compare models and estimate parameters. Our three summary statistics were: 1) the difference in mean phenotype between 2011 to 2015 (which we measured in the resurrection experiment), 2) the difference in mean phenotype between 2015 and 2017 (measured in the resurrection experiment), and 3) the ratio of survival rates between the drought (averaging 2012-2015) and pre-drought (averaging 2006-2011) periods (measured in the field). The observed values of the first two summary statistics were calculated using the estimated marginal means from the linear mixed effects models described above in *Statistical analyses*. For the observed value of the third summary statistic, because survival in the model is only affected by the focal trait but in the field is likely affected by multiple traits, we assumed that one third of the decrease in survival during drought in natural populations was due to the focal trait (flowering phenology).

**Table 1.** Description and parameter prior distributions for the four candidate models.

|  | Description | Parameters |  |  |
| --- | --- | --- | --- | --- |
|  |  | Seed survival rate | Seed germination rate | Change in optimum |
| **Model A** | No seed bank, with the trait optimum changing at the beginning of the drought, then returning to the historical optimum after drought | 0.0 | 1.0 | U[-20.0, 0.0] |
| **Model B** | No seed bank, with the trait optimum changing at the beginning of the drought and staying the same post-drought | 0.0 | 1.0 | U[-20.0, 0.0] |
| **Model C** | With seed bank, with the trait optimum changing at the beginning of the drought, then returning to the historical optimum after drought | U[0.05, 0.75] | U[0.05, 0.75] | U[-20.0, 0.0] |
| **Model D** | With seed bank, with the trait optimum changing at the beginning of the drought and staying the same post-drought | U[0.05, 0.75] | U[0.05, 0.75] | U[-20.0, 0.0] |

For each of our two populations, we explored four models (Table 1). Models were run separately for each population because model demography was explicitly tied to observed census sizes in each population (see above). In the models without seed banks (Models A and B), seed survival was set to 0.0 and seed germination was set to 1.0, while the change in optimum was allowed to vary between −20.0 and 0.0 (Table 1). For models with seed banks (Models C and D), seed survival and germination rates were also allowed to vary between 0.05 and 0.75, which encompasses the range of values for these parameters estimated across 20 *C. x. xantiana* populations in [42] (range of posterior mode for germination = [0.11, 0.27]; seed survival [0.45, 0.72]). We ran 20,000 simulations of each model for model comparison, then ran an additional 20,000 simulations of the best model to aid in parameter estimation. We used the abc package [63] in R to compare models and estimate parameters (*Supp. Mat. D*). Graphical posterior predictive checks were performed to compare observed summary statistics in each population to the distribution of summary statistics resulting from 500 simulations run using parameter values drawn from the estimated parameter posterior distributions.

To estimate the individual effects of aboveground demography (seed input), seed bank vital rates (survival and germination), and selection environment (change in phenotypic optimum during drought) on the evolutionary stasis of KYE relative to S22 (Results), we compared the posterior predictive checks described above to additional batches of simulations (500 simulation each) as follows:

1. Aboveground demographic effect: seed bank (survival and germination) and optimum change parameters estimated for KYE, but with demographic history of S22
2. Seed bank effect: optimum change estimated for KYE, with KYE demographic history, but with seed bank parameters estimated for S22
3. Selection environment effect: seed bank parameters estimated for KYE, with KYE demographic history, but with optimum change parameter estimated for S22

## Results

### Climate and demography

Both sites experienced a period of substantially reduced precipitation in years 2012-2015 (Fig 1). For KYE, growing season precipitation was reduced by 45% compared to 2006-2011; for S22, precipitation was reduced 57%. Precipitation increased in 2016 and 2017 at both sites, with particularly high precipitation in 2017.

**Figure 1.**
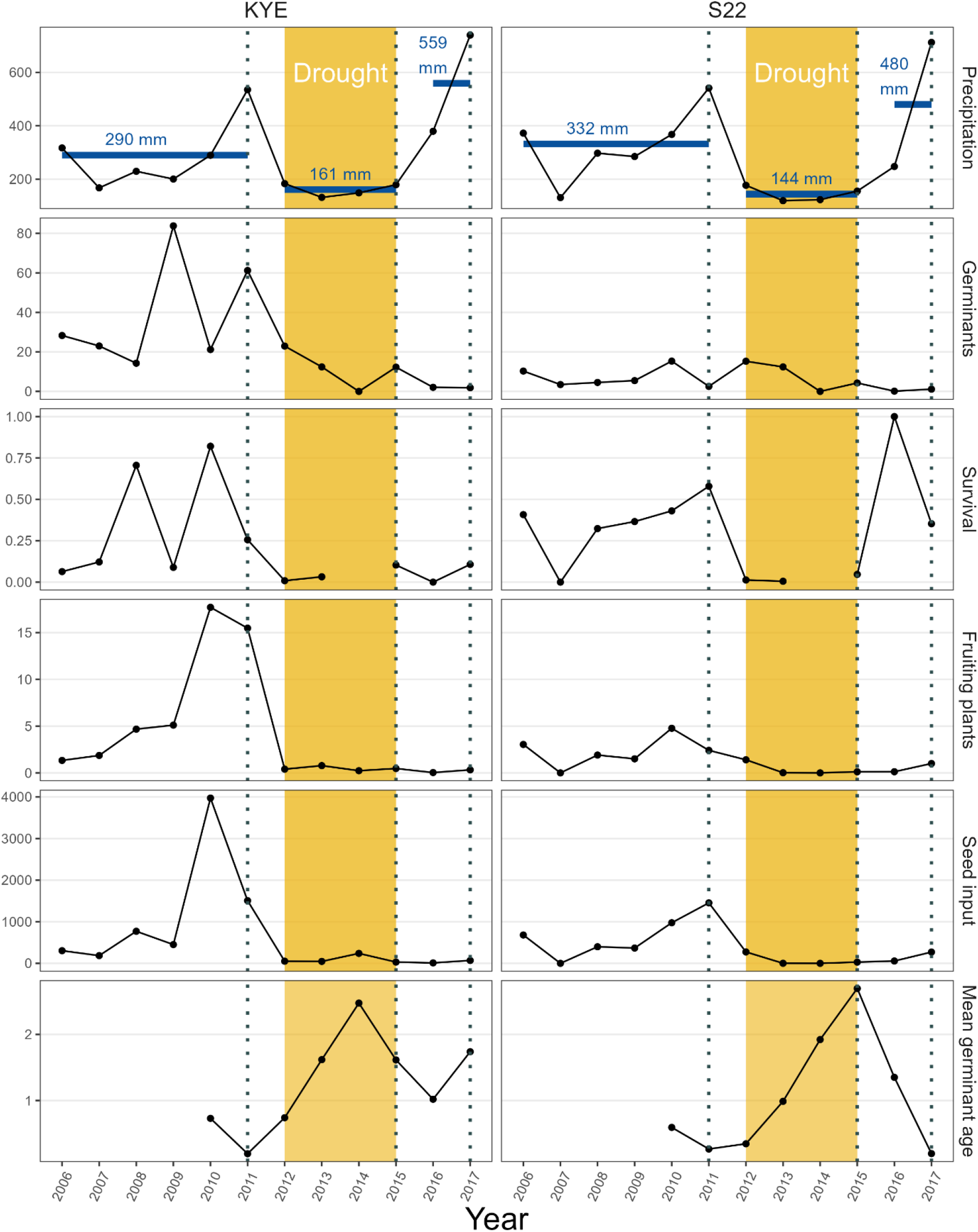
Environmental and demographic trends in two populations (KYE, S22) of the California annual plant *Clarkia xantiana ssp. xantiana* from 2006-2017. Dotted vertical lines mark cohorts collected for the resurrection experiment. Precipitation represents the cumulative growing season precipitation, in mm, for a given year (i.e., precipitation for year 2011 was the cumulative precipitation from November 2010 – June 2011). Blue horizontal bars in top panels mark the mean precipitation across three periods – pre-drought, during drought, and post-drought, with the drought period (2012-2015) highlighted in gold. Germinants, fruiting plants, and seed input are per m^2^. Survival rate is based on censuses of germinants and fruiting plants in permanent plots in each population; no survival data are presented for 2014 because zero germinants were recorded in both populations. Mean germinant age in each year is estimated based on seed input in years prior and seed bank vital rates from [42].

Overall, KYE, which is nearer the range center, had higher density and total population size than S22, which is located near the range edge (average density across 12 years: 4.03 and 1.36 fruiting plants per m^2^, respectively; average population size: 282,704 and 9,166 fruiting plants). Reduced precipitation during the 2012-2015 drought period had strong demographic effects at both sites (Fig. 1). At both sites, survival rates of plants in permanent plots decreased over 85 percent during drought relative to pre-drought averages (KYE survival rates: pre-drought = 0.34, drought = 0.05; S22 pre-drought = 0.35, drought = 0.02). At KYE, fruiting plant estimates during this drought period were 94% lower than pre-drought estimates (0.5 and 7.7 fruiting plants per m^2^, respectively), though the pre-drought average was largely influenced by high abundances in 2010 and 2011 (Fig. 1). At S22, plant densities during the drought period were 83% lower than pre-drought estimates (0.4 and 2.3 fruiting plants per m^2^, respectively). In the two years post-drought, fruiting plants remained few at KYE. At S22, the number of fruiting plants somewhat rebounded in 2017.

Seed input prior to the drought was roughly twice as high at KYE relative to S22 (yearly mean of 1,198 and 646 seeds per m^2^, respectively). Seed input was especially high at KYE in the two years preceding the drought, 2010-2011. Seed input was similar at KYE and S22 during the 2012-2015 drought period (92 and 76 seeds per m^2^, respectively; Fig. 1). In the two years post-drought, patterns of seed input mirrored patterns of plant density, with input remaining low at KYE but trending upward at S22 in 2017. The decreased germination and increased seed survival rates of KYE (estimated in [42]) along with high seed input resulted in KYE having a more substantial seed bank than S22, with estimated mean germinant age ca. 20% higher than S22 (1.27 vs. 1.04 mean germinant age, respectively; Fig. 1). In both populations, low seed input during drought resulted in increased germinant age during that period due to older seeds comprising a larger proportion of the seed bank (Figs. 1, S8).

### Phenotypic evolution

For S22, days to flowering differed among year cohorts (*P* = 0.004; Fig. 2; Table S3). Days to flowering decreased 1.6 days from 2011 to 2015, and another 1.2 days from 2015 to 2017. Pairwise Tukey tests identified the 2011 / 2017 contrast in days to flowering as significant (69.4 vs. 66.4 days, respectively; *P* = 0.006). Phenotypic variance in days to flowering also decreased across year cohorts, from 28.6 in 2011, to 21.3 in 2015, to 14.3 in 2017 (Table S2). There was also a trend in S22, albeit not significant, for growth rate to increase over this timeline, which may have contributed to earlier flowering time. There was no statistically significant evidence of phenotypic evolution of other traits for S22, nor any traits in the KYE population.

**Figure 2.**
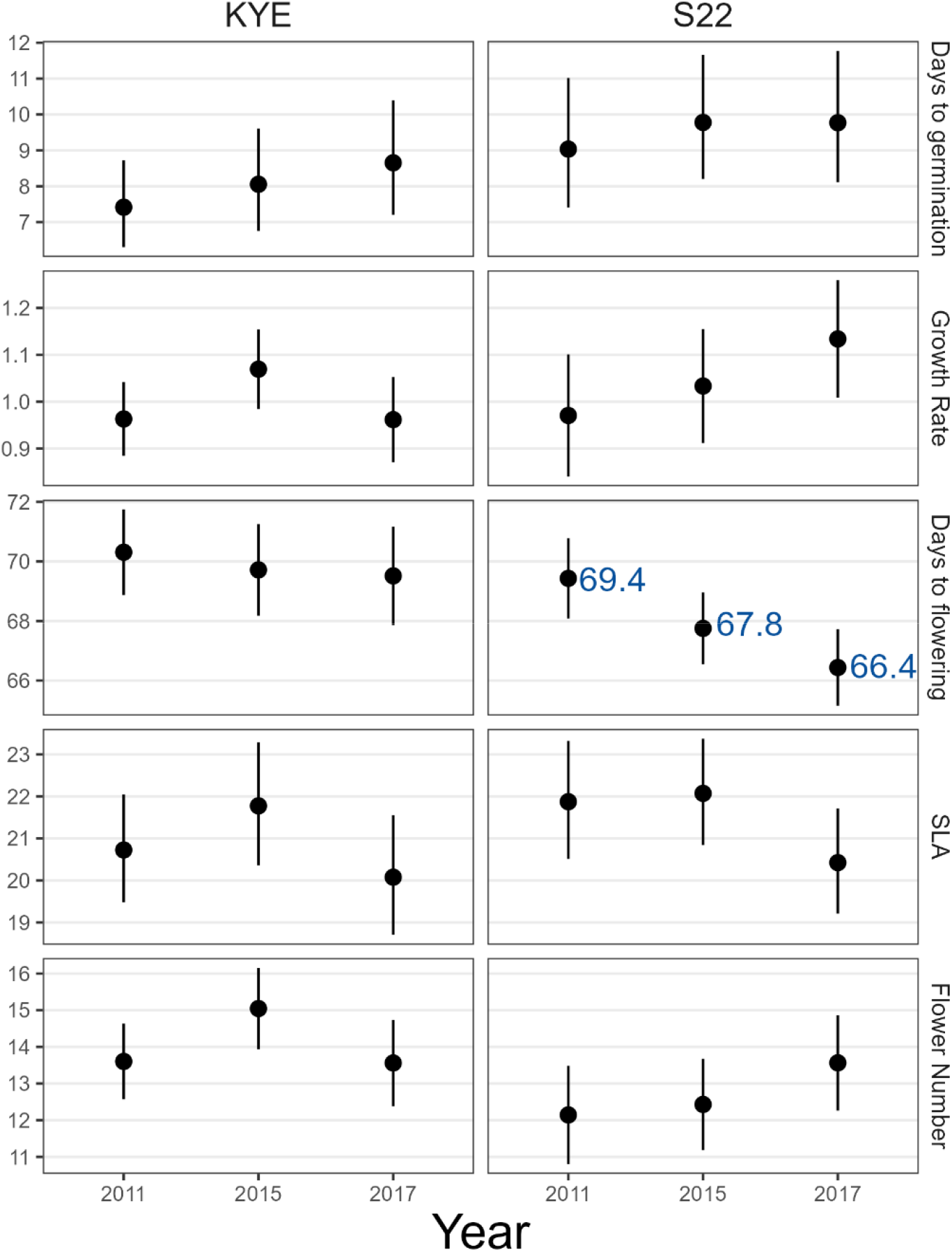
Measured phenotypes across the three sampling years for populations KYE and S22. Each panel shows estimated marginal means with 95% CI’s estimated from the model *trait ~ year + transplant age + (l/sire/dam)*. Days to germination was measured as days elapsed between sowing and emergence; growth rate was measured as leaves produced per day; days to flowering was measured as days elapsed between sowing and opening of the first flower; SLA was calculated as mm^2^ mg^-1^; flower number was measured as the total number of flowers an individual produced in its lifetime. Raw data are shown in Fig. S6.

### Additive genetic variance

There was evidence of significant additive genetic variance () for days to germination, flowering date, and flower number in both populations (Fig. 3; Table S4) The *h*^2^ estimates for these traits ranged from 0.32 – 0.59. There was also evidence of significant for growth rate in the S22 population, but not in the KYE population. We did not find evidence that for SLA was significantly different from zero in either population.

**Figure 3.**
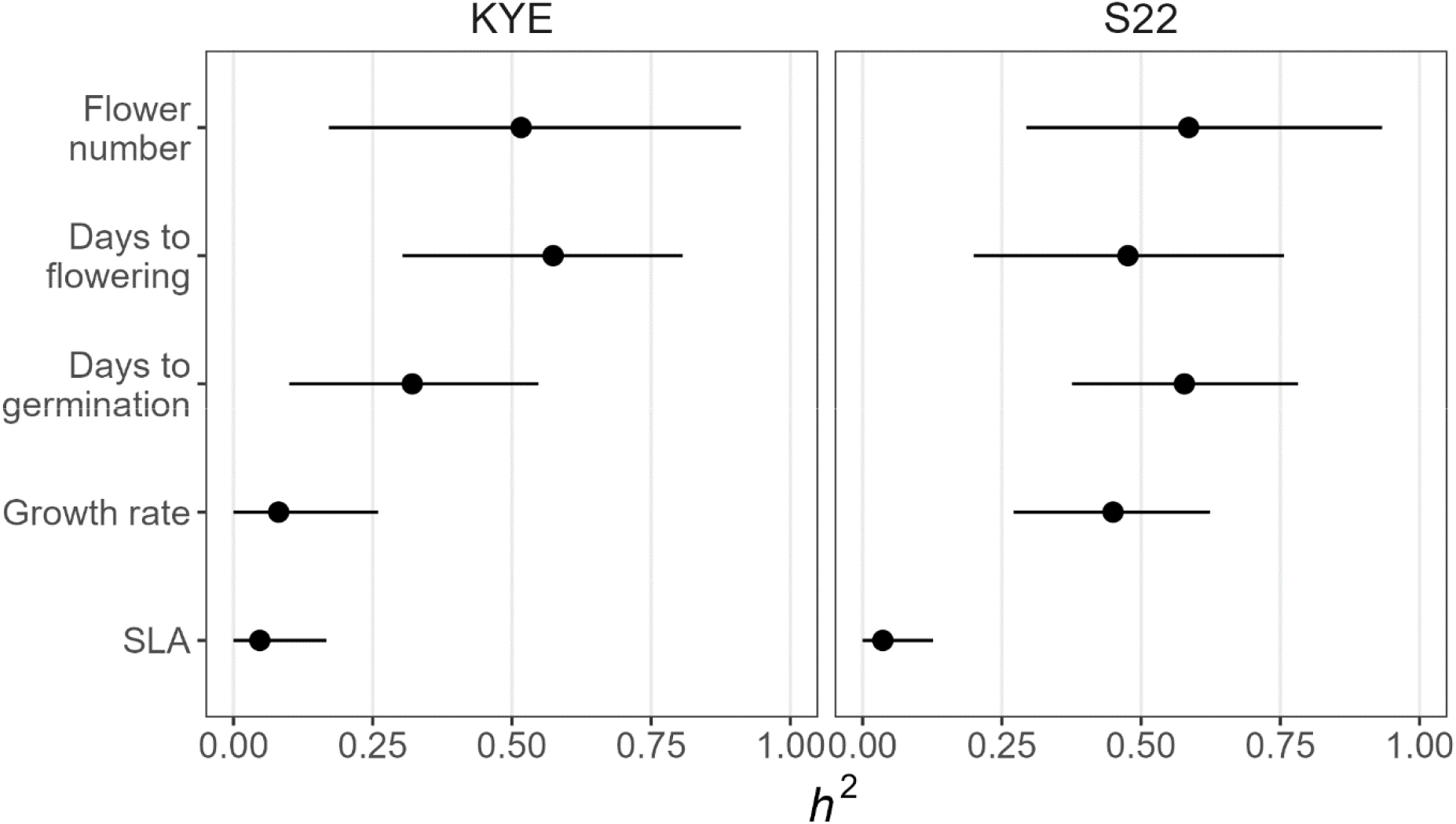
Estimates of narrow-sense heritability (*h*^2^) for traits, with 95% credible intervals. Estimates of additive genetic variance for each trait are in Table S4.

### Simulation model

For both populations, the model with the highest posterior probability was Model D (population with a seed bank, with post-drought phenotypic optimum equal to optimum during drought) (Table S5). Posterior probability of Model D was much higher than all other models for KYE (Bayes factors all > 12; Table S5). For S22, the contrast of Model D with the second best model, Model B (population *without* a seed bank, with post-drought phenotypic optimum equal to optimum during drought), was weaker (Bayes factor = 2.3), indicating less support for an influential seed bank in that population compared to KYE. Overall, without a seed bank, populations experiencing a strong demographic decline due to large changes in phenotypic optima saw rapid phenotypic evolution toward the new optimum (Fig. S9a). However, with a seed bank, even large changes in optima usually resulted in only modest phenotypic evolution like that observed in our populations. These constrained evolutionary responses were caused by gene flow from prior generations via the seed bank.

We can concisely describe the strength of a population’s seed bank as a combination of the seed germination and survival rates, *ϕ* = (1 – *g*)*s*. Thus, *ϕ* (“strength of seed bank” in Fig. 4) describes the expected proportion of seeds available for germination in year t that will be available for germination in year *t* + 1, with larger values of *ϕ* indicating a stronger seed bank. The strength of the seed bank at KYE was estimated to be more than five-fold higher than that of S22 (0.47 vs. 0.07) due to a higher seed survival rate and a lower germination rate (Fig. 4a). KYE was also estimated to have experienced a 15% smaller change in optimum during drought compared to S22 (−6.1 vs. −7.2 days; Fig. 4a). Observed evolutionary and demographic responses of both populations fell within the distributions of posterior predictive checks (Fig. 4b).

**Figure 4.**
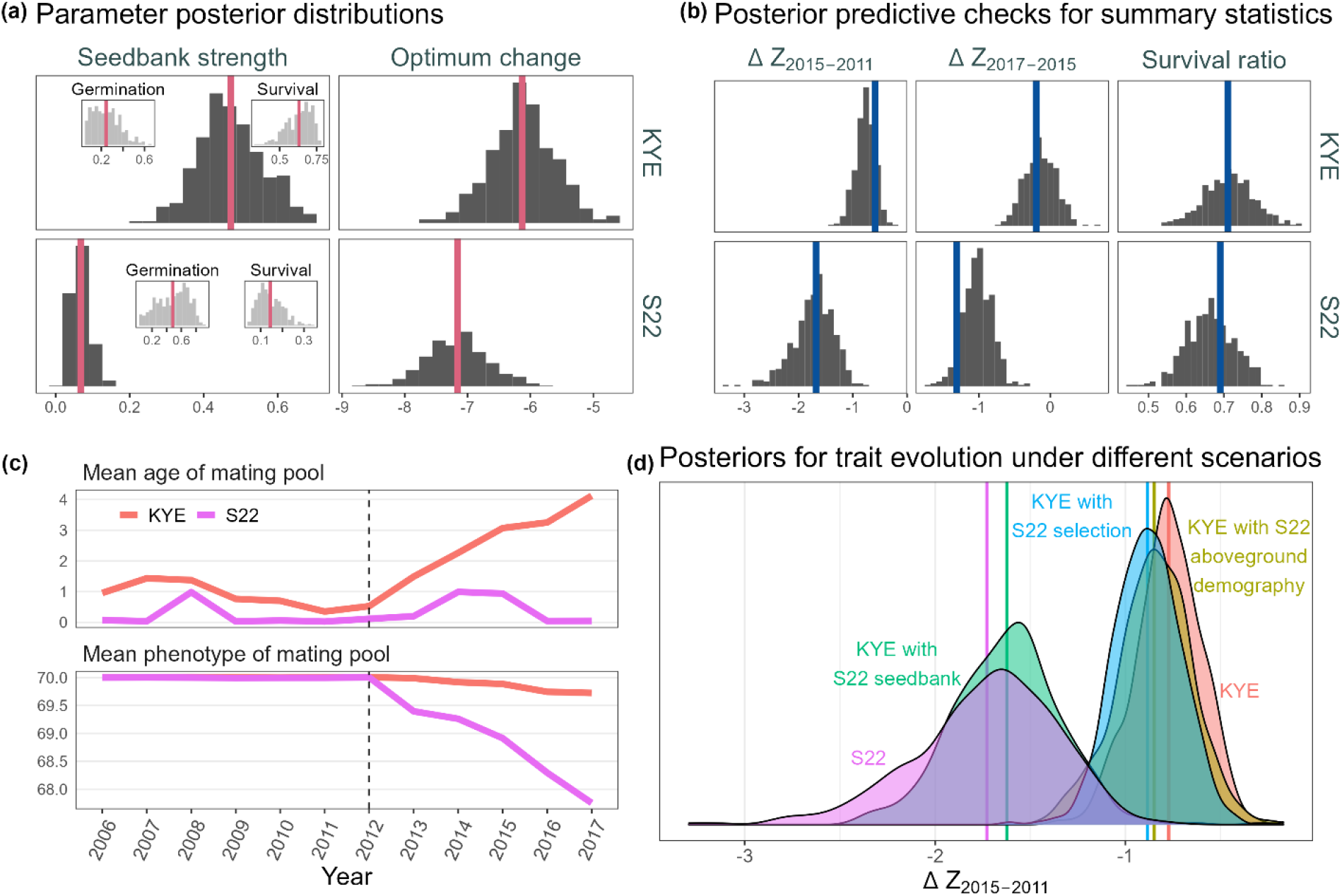
Results of evolutionary simulations and ABC analyses. (a) Estimated posterior distributions for model parameters (germination rate, seed survival rate, change in trait optimum during drought). Seedbank strength () is a joint parameter defined as, where *g* is germination rate and *s* is seed survival rate. Red vertical lines mark the posterior means. (b) Posterior predictive checks for the three summary statistics used in ABC analyses (change in trait mean 2015-2011, change in trait mean 2017-2015, and ratio of survival during drought to survival pre-drought). Blue vertical lines mark observed summary statistics from empirical data. (c) Mean age and phenotype of mating pools in both populations over the 12 focal years during the simulation. Dashed vertical line marks the year when the trait optimum changed. (d) Distributions of the summary statistic (change in trait mean 2015-2011) under varying model scenarios. Colored vertical lines mark posterior means for each scenario.

KYE’s mating population had a consistently higher mean age than S22’s because of its stronger seed bank (Fig. 4c). However, mean age for both populations increased during the drought period due to low fecundity and increased representation of older generations. The increased gene flow from earlier generations in KYE led to the mean phenotype of its mating pool responding much more slowly to the environmental perturbation than S22 (Figs. 4c, S9b). Further simulation runs indicated that the difference in evolutionary responses was overwhelmingly due to differences in seed bank vital rates, as opposed to differences in aboveground demography or selective environments (Fig. 4d). Simulations run with the selective environment and aboveground demography of KYE, but with the seed bank rates of S22, produced evolutionary change nearly equal to that observed in simulations run with full S22 parameters.

## Discussion

Extreme environmental perturbations offer an opportunity to examine rapid evolution in natural populations. We used a resurrection experiment to test whether native plant populations rapidly evolved in response to a multi-year drought despite severe demographic declines. In one population (S22), earlier flowering evolved by the end of the drought and continued to evolve in the same direction after the drought ended. However, no other traits diverged at S22, and all traits exhibited stasis in the second population, KYE. We detected substantial additive genetic variance for multiple traits in both populations, including flowering phenology, suggesting that responses to selection were not constrained by limited genetic variation. Evolutionary simulations integrated with our demographic and experimental data suggested that seed banks constrained evolutionary responses in both populations. In KYE, the seed bank had a stronger effect resulting in evolutionary stasis; whereas in S22, the seed bank had a weaker effect, which facilitated closer tracking of changing phenotypic optima.

Why does rapid evolution occur in some populations but not others? One hypothesis is that a lack of quantitative genetic variation limits adaptive responses even when selection is intense. However, we found substantial *V_A_* and high heritability for flowering phenology in both populations. In fact, KYE contained twice the additive genetic variance in flowering time compared to S22. Thus, these results are inconsistent with the hypothesis that genetic variation prevented rapid evolution. Other studies have similarly found that rapid evolution occurred in some but not all studied populations [12–14]. However, estimates of *V_A_* are rarely quantified in this context and thus the importance of genetic constraints is poorly understood [but see 5]. It is also worth noting that phenotypic variance in flowering phenology in S22 decreased 50% over our study, consistent with a strong episode of selection. Thus, the erosion of genetic variation due to this drought could constrain future responses to selection on phenology.

A more likely explanation for the lack of phenological evolution in KYE is that gene flow from the seed bank reintroduced alleles that were maladaptive during the drought. Simulations suggested that decreased germination and increased seed survival rates resulted in a seed bank at KYE that had a stronger effect on responses to selection compared to S22. The recruitment and mating of older individuals from the seed bank essentially caused maladaptive gene flow through time (Fig. 4c). Similarly, a third population grown during the refresher generation showed no significant evolutionary response to the drought (Fig. S4) and was estimated in [42] to have seed bank vital rates similar to KYE (low germination, high seed survival). It would be fruitful for future resurrection studies to explicitly sample populations across a range of seed bank vital rates to further dissect seed bank effects on adaptation to climate-change-induced selective episodes. Additionally, seed banks can serve to buffer populations from extinction during times of environmental stress [64,65], and the interplay between these two contrasting consequences of seed banks for populations is an area ripe for study [22,28].

Of course, “all models are wrong” [66], and our simulation models do not perfectly reflect reality for these natural populations. But they do demonstrate that, for a population with substantial genetic variance for a trait under selection during an environmental perturbation with severe demographic consequences, like our populations during the megadrought, selection is expected to effectively drive rapid phenotypic evolution toward new trait optima. Seed banks can greatly constrain this evolution and produce results in line with our own empirical data, with parameter estimates that follow patterns of empirically estimated seed vital rates. However, the hypotheses tested above are not exhaustive, and it is worth exploring alternative explanations for the lack of evolutionary change in phenology at KYE. Trait plasticity could allow a population to weather an environmental perturbation with little or no evolutionary change [67]. However, plastic responses in flowering time were nearly identical for our two populations (Fig. S10), suggesting that greater plasticity in KYE cannot explain its evolutionary stasis. Spatial gene flow among populations could potentially oppose allele frequency changes favored by selection. However, gene flow is unlikely to be strong enough over this short time window to prevent evolutionary change, especially given that our populations are isolated from others (>1 kilometer) and lack adaptations for long-distance dispersal. It is also possible that advanced phenology simply did not confer fitness benefits during drought at KYE, though this would be contrary to both general trends in plants [44] and the well-documented relationship between early flowering time and adaptation to aridity in *C. xantiana* and close relatives [37,45,61,68–70]. It is also interesting that, in contrast to expectations, empirical and simulation results suggested that phenotypic optima did not return to their pre-drought level after the drought ended. This insight is consistent with the observation that population demography did not rebound after drought as one might expect (Fig. 1). Although the causes of this trend are unclear, it could be related to the timing of precipitation [e.g., most precipitation fell early in the season, which seems to promote germination but not survival in *C. x. xantiana* (DM, JB, pers. obs.)].

The evolution of earlier flowering at S22 is consistent with post-drought resurrection studies in *Brassica rapa* [5,33], and likely reflects a strategy of drought escape [reviewed in 44,71,72]. Past work on *C. x. xantiana* has shown that Q_ST_ for flowering phenology is more than twice as high (Q_ST_ = 0.7) than Q_ST_ for five other ecologically-important traits [68]. This result suggests that flowering phenology is often the target of spatially-variable selection and readily evolves, as has been frequently observed in many systems [5,44,73,74]. Earlier flowering may evolve more readily than other traits under selection if its genetic architecture is comparatively simple [e.g., 71], which can quicken responses to selection relative to more highly polygenic traits. The evolutionary lability of flowering phenology could also be due to its direct tie to assortative mating [75] — early flowering individuals tend to mate with other early flowering individuals. Theoretical work has shown that when there is directional selection on flowering time, positive assortative mating can increase genetic variation and the rate of phenotypic evolution compared to scenarios with random mating [76,77].

A population’s response to environmental change will be determined by the interplay of demography, genetics, selection, and stochastic processes. Here, we have shown that flowering phenology rapidly evolved during and after a severe drought in one population. Whereas, in a second population we observed no sign of rapid evolution despite similar environmental stress, demographic decline, and even greater additive genetic variance in phenology. In both populations, our results are consistent with the hypothesis that gene flow through time via seed banks either slowed or prevented rapid evolution. Our results are inconsistent with the hypotheses that plasticity or limited genetic variation constrained responses to selection. While rapid evolution in response to environmental perturbations has been repeatedly demonstrated, few studies have tackled the problem of why evolutionary stasis is so commonly observed across studies and systems. Our results emphasize that studies of adaptation in plant populations experiencing environmental change should consider both above and belowground processes affecting population demography and fitness. As we seek to build more predictive models of the evolutionary process, the synthesis of demographic, environmental, and quantitative genetic data will be invaluable for understanding where and when rapid evolution is likely to occur.

## Supporting information

Supp. mat.

## Acknowledgments

We thank Monica Geber and Vince Eckhart for their assistance with collection of seeds, demographic and environmental data, and for their comments on a previous version of this manuscript. We thank Eric Bakken, Ryan Allen, Hannah Littel, Samantha Sorg, and Adam Kostanecki for assistance with experiments. Conversations with Ruth Shaw and Seema Sheth were invaluable in the design of the experiments.

