## Supplementary material for "Rapid evolution in response to climate-change-induced drought and its demographic and genetic controls": Supp. mat.

### A: Demographic data

Demographic data were collected as part of a long-term study of 36 populations across the range of *C. x. xantiana* in years 2006-2017. Each winter (January / February), germinants were counted in 20-30 long-term, 0.5 m2 permanent plots; these same plots were censused for fruiting plants in summer, allowing us to estimate survival rate as the number of fruiting plants divided by the number of germinants. We were unable to calculate survival rates in 2014, as there were zero germinants recorded in permanent plots at both sites (though one fruiting plant was recorded at KYE). As a proxy for population size, we obtained counts of the number of fruiting plants at each site, as estimated from censusing 58-129 0.5 $m^2$ quadrats across the extent of each site in late June / early July of each year. We report these data as the average number of fruiting plants per $m^2$, and obtain a rough estimate of total population size by multiplying this number by the total area of the site (KYE: 7.015 ha; S22: 0.674 ha). We also recorded the number of fruits per plant on a subset of these plants (mean of 132 plants per site per year). We obtained estimates of the number of seeds per fruit from fruits collected from across each population (mean of 27 fruits per site per year). We report the estimated seed input per plot as the product of the average number of fruiting plants per plot, the average number of fruits per plant, and the average number of seeds per fruit.

#### Seed bank estimation

We combined our observed data on seed input for each population in each year with estimates of seed survival and germination to estimate the proportional representation of year cohorts among germinants and calculate mean germinant age in each year. Seed input for each population in each year was estimated as described in the main text (*Materials and Methods: Climatic and demographic data*). We used estimates of germination and seed survival rates (the mode of each rate's posterior distribution) from [1], which are based on multiple years of field trials. In *C. x. xantiana*, seeds are produced in July and germinate in January / February. For our purposes, seed survival ($s$) was calculated as the product of $s_2$ (survival of ungerminated seeds from January/February to the next October) and $s_3$ (survival from October to the next germination opportunity in January/February) in [1]. We assumed germination rate ($g$) was the same for all seed ages, as only one germination rate was estimated for each population in [1]. These rates correspond to $s$ and $g$ used in the simulation models, described below in *C: SLiM model*. We assumed $s_0$ and $s_1$, which in [1] represent seed survival between initial seed production and the first germination opportunity the following January / February, were both equal to 1 — i.e., all seeds produced survived to the first germination opportunity. This does not influence the relative proportion of different seed cohorts in the seed bank (as all cohorts suffer that initial seed death at the same rate).

Representation of each cohort in the germinant pool each year was calculated using seed input ($i$), $s$, and $g$. We assumed seeds did not survive beyond three years so that we could calculate mean germinate age for an appreciable number of years (e.g., if we assumed seeds survived five years, we could not calculate mean germinant age for any pre-drought years, as our demographic data only extends to 2006). Thus for the germinant pool in year *t*, cohort representation (germinant ages 0 - 3) was calculated as,

$$germinants_0 = i_{t-1}g,$$
$$germinants_1 = i_{t-2}(1 - g)sg,$$
$$germinants_2 = i_{t-3}(1 - g)s(1 - g)sg,$$
$$germinants_3 = i_{t-4}(1 - g)s(1 - g)s(1 - g)sg$$

See archived R code for full workflow.

### B: Invisible fraction bias

Invisible fraction bias can occur in resurrection experiments if non-random mortality of propagules during storage biases the estimation of a population's mean phenotype, due to genetic correlations between these survival traits and other traits of interest [2]. The recommended method to test for invisible fraction bias is to calculate the correlation between proportion germination and the traits of interest for maternal families. These data are usually collected for the refresher generation. In our refresher generation, we sowed two seeds per planting tray cell and culled one germinant if two germinants emerged, but did not record the number of germinants in each cell. Thus, we did not have the exact data needed to test for the invisible fraction bias as recommended in [2].

Instead, we used an alternative approach to ask whether there is likely a bias in our estimated population means due to unequal germination rates among maternal families in the refresher generation. Taking the refresher generation data, we plotted trait means for each population / year cohort based on two different calculations: first, the raw, observed data for all individuals (blue points in Fig. S1 below), with varying number of individuals from different maternal families; second, a "corrected" mean (red points in Fig. S1) that assumes equal representation for all maternal families (by averaging across family means). As shown in Figure S1, there did not appear to be any substantial biases in our estimations of mean trait values in the refresher generation due to unequal germination among maternal families.

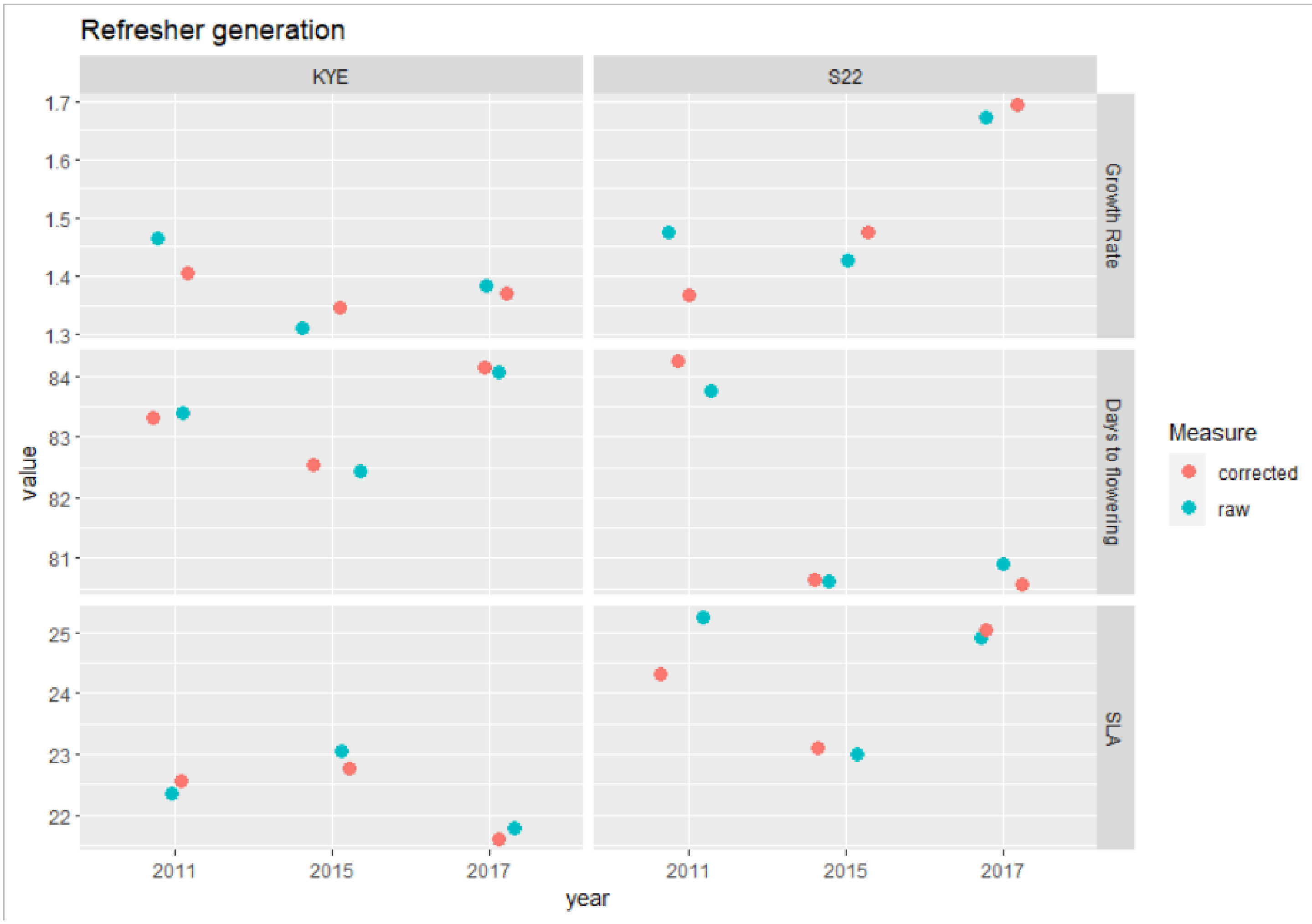


**Figure S1.** Mean trait values for each population / year cohort in the refresher generation, colored by whether the mean was calculated based on raw data or as the average of family means ("corrected").

For the measurement generation, we did record germination data for individual seeds (summarized in Table S1 below). Thus, in contrast to the refresher generation, we were able to calculate the correlation between germination proportion and our focal traits across maternal families. Correlations were weak to moderate (max $|r| = 0.33$; Fig. S2). Using the "correction" approach as in the refresher generation above, Figure S3 shows that there do not appear to be any large biases in our estimations of mean trait values in the measurement generation due to unequal germination among maternal families.

**Table S1.** Proportion germination for each population / year cohort in the measurement generation.

| | **KYE** | **S22** |
|---|---|---|
| **2011** | 0.73 | 0.63 |

| | | |
|---|---|---|
| **2015** | 0.69 | 0.75 |
| **2017** | 0.63 | 0.61 |

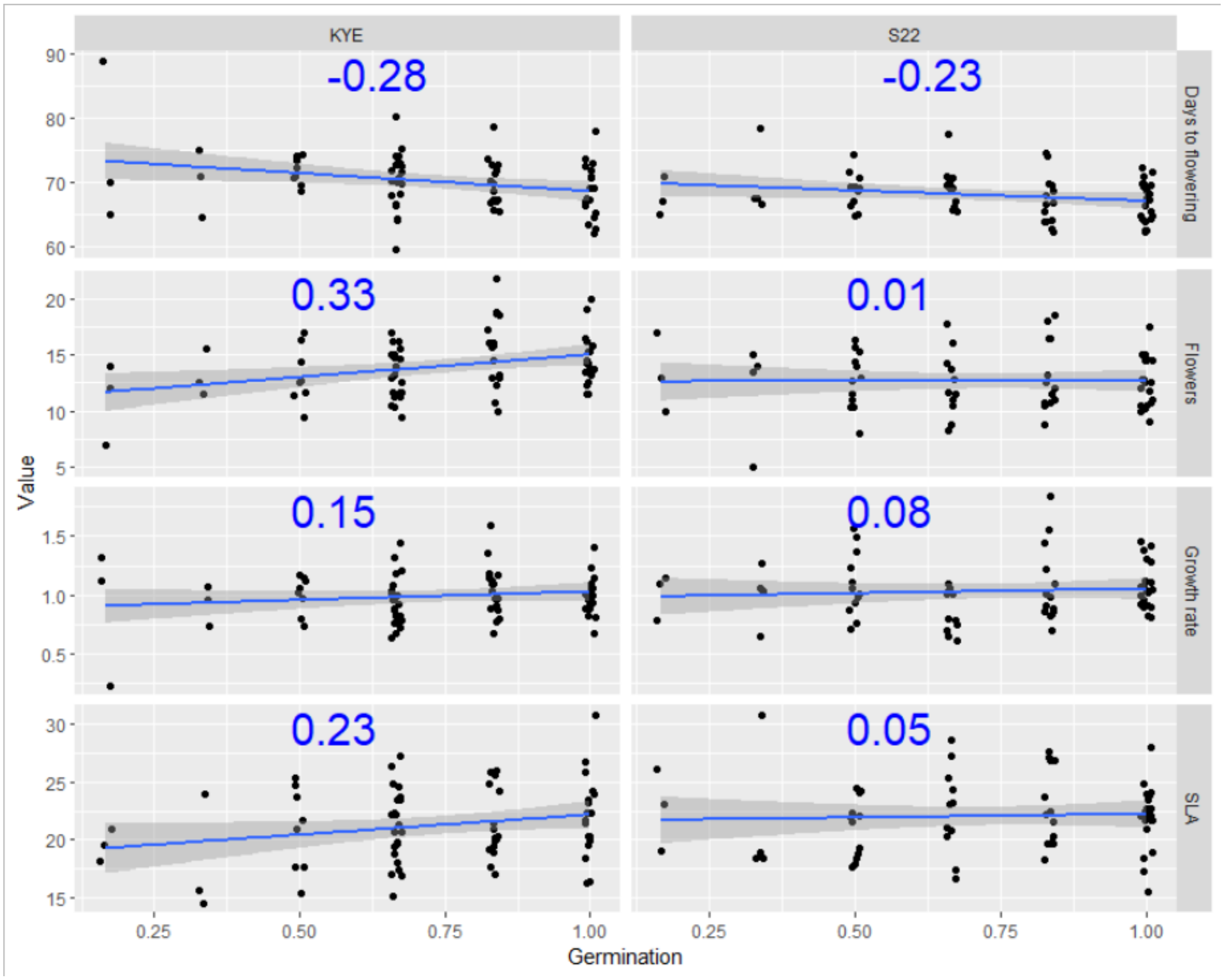


**Figure S2.** Relationship between maternal family proportion germination (X-axis) and focal trait means (Y-axis). Pearson correlation coefficients between maternal family germination proportion and focal traits in the measurement generation are shown in blue text. Blue line is linear regression of trait ~ germination.

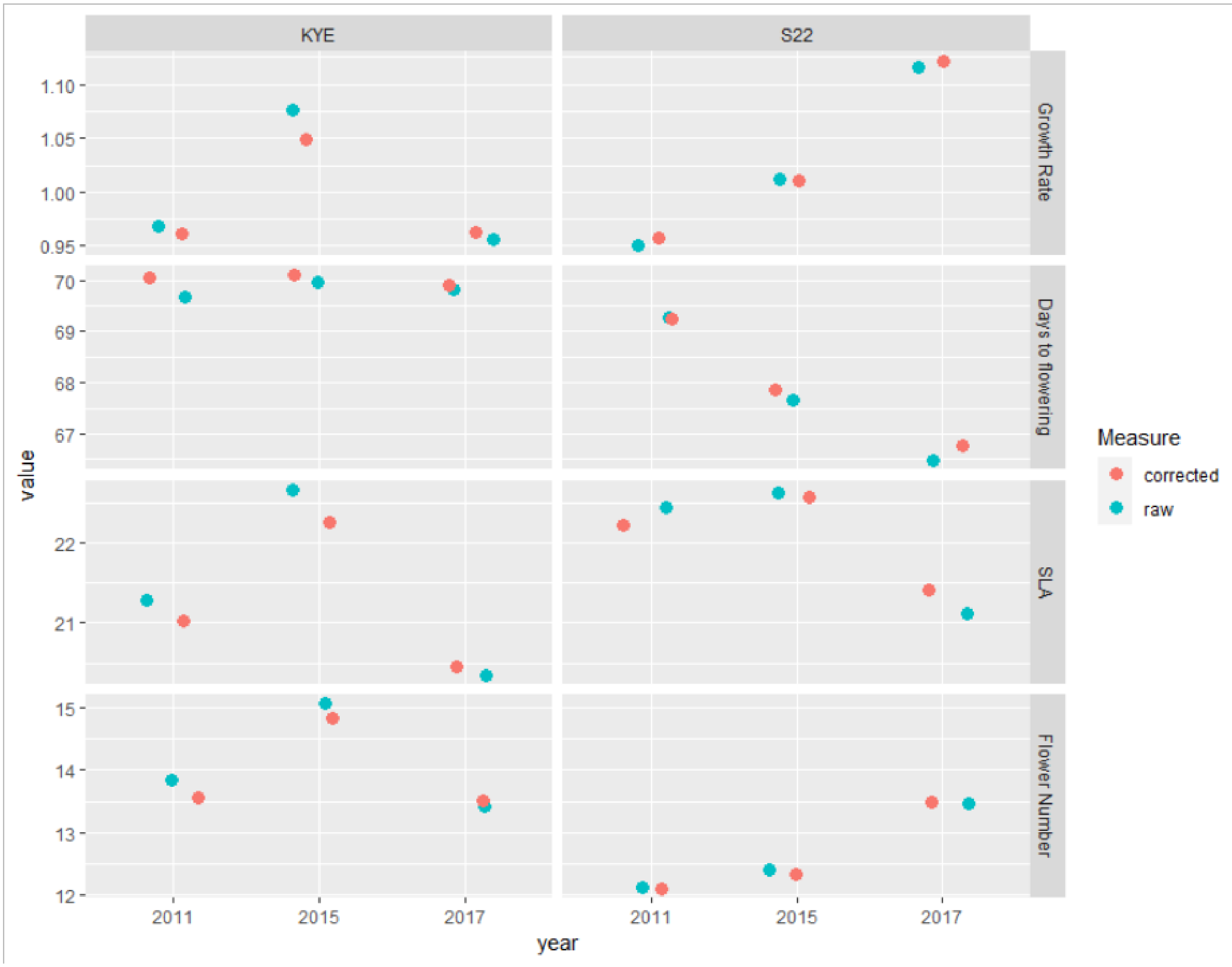


**Figure S3.** Mean trait values for each population / year cohort in the measurement generation, colored by whether the mean was calculated based on raw data or as the average of family means ("corrected").

### C: Quantitative genetic analyses

We estimated quantitative genetic variance components for all traits in both populations using an animal model implemented in the MCMCglmm package. Days to germination and SLA were log transformed prior to analysis. Each model included year and transplant age as fixed effects (except for the model of days to germination, where we used dam seed weight instead of transplant age) and animal as a random effect (in MCMCglmm, individuals are "animals" and this model form is used to estimate additive genetic variance). We used weakly informative priors for each model: an inverse-Gamma prior for residual variance (V=1, nu=1) and a parameter-expanded prior for the random effect (V=1, nu=1, alpha.mu=0, alpha.V=1000). We ran each model for 800,000 iterations with a burn-in of 15,000 and thinning of 250. The effective sampling sizes for random effects ranged from 2,038-3,442. To test for maternal environmental effects, we also built models with *dam* as a random effect in addition to *animal* and compared these models to the model without *dam* using deviance information criterion (DIC). If including the *dam* term improved model fit (i.e., smaller DIC value), we used this model including maternal environmental effects to estimate variance components [3]. For KYE, growth rate and SLA models included a dam term. We used parameter-expanded priors because these often result in better mixing of the MCMC chains and allow more flexibility in the shape of the prior [4]. Using a weakly informative prior without parameter expansion (V=1, nu = 0.002 for both residual variance and random effects) resulted in qualitatively similar patterns in $h^2$ overall, though estimates of $h^2$ for days to germination and growth rate for the S22 population, and growth rate and SLA for the KYE population, were higher compared to results using parameter expanded priors (Fig. S7).

### D: SLiM model

We used an individual-based simulation model paired with Approximate Bayesian Computation (ABC) to test hypotheses about the influence of seed banks and selection environments on the evolutionary dynamics of flowering time in our two populations. Using SLiM [5], a forward-in-time population genetic modeling software, we built an individual-based simulation of the evolutionary and demographic dynamics of a population experiencing an abrupt environmental perturbation. Fitness was based on the deviation of an individual's phenotype from the phenotypic optimum, which varied through time. The population experienced a burn-in period of 1,000 generations with a constant carrying capacity ($K$ = 10,000) during which mutation, recombination, and gene flow increased genetic variance to levels comparable to those observed in natural populations. Following the burn-in, the model simulated the focal 12 years captured in our long-term demographic data, with carrying capacity proportional to census size recorded during those years so as to simulate observed population demographic patterns. The phenotypic optimum decreased during the years corresponding to drought in our natural populations, and then either returned to the pre-drought optimum after the drought ended, or stayed at the drought optimum, depending on the model scenario (see *Simulation model* in main text). SLiM code is archived alongside the other data and analyses associated with this manuscript.

#### Genetics and mutation

In our simulations, individuals were diploid and hermaphroditic, with obligate sexual reproduction, and a single chromosome 100,000 bp long. We modeled one mutation type that contributed additively to the focal quantitative trait [i.e., biallelic quantitative trait loci (QTL) with no dominance]. The overall mutation rate (mutations per base position on a gamete per generation) was set to $1 \times 10^{-8}$. QTL effect size was drawn from a standard normal distribution [i.e. N(0, 0.25)]. Recombination rate (probability of a cross-over event between any two adjacent bases per genome per generation) was set to $1 \times 10^{-5}$. Narrow-sense heritability of the focal trait was set to $h^2 = 0.5$.

#### Fitness, mating, and population dynamics

Female viability ($W_i$), the probability of surviving to set fruit, was under stabilizing selection and, as in [6], calculated as

$$W_i = exp[-\frac{(z_i - \theta_t)^2}{2v^2}] \quad (1)$$

where $v$ is the strength of selection (the standard deviation of the stabilizing fitness function), and $z_i - \theta_t$ is the deviation of individual $i$'s trait value ($z_i$) from the phenotypic optimum at time $t$ ($\theta_t$). We set $v = 5.27$, which was the average observed phenotypic standard deviation in days to flowering among the six population / year cohorts. [To illustrate this fitness function quantitatively: an individual with a phenotypic deviation (from the optimum) of 2 would have P(fruiting) = 0.88; an individual with a deviation of 5 would have P(fruiting) = 0.46.] The fruiting status for an individual (did or did not set fruit) was determined by a draw from a binomial distribution with probability equal to $W_i$. Individual fecundity was drawn from a Poisson distribution with mean ($f$) = 20. The father for each offspring (seed) was drawn at random (with replacement) from the population; siring success was not influenced by an individual's phenotype. Populations were subject to negative density dependence based on carrying capacity ($K$); K was set equal to 10,000 during burn-in and fluctuated proportional to observed census sizes during the focal 12-year period (see *Simulation Process* below).

For scenarios with a seed bank (models C and D), one extra patch acted as the seed bank. Each generation, all offspring initially migrated to the seed bank. A fraction of the individuals in the seedbank (which contained new seeds as well as older seeds) "germinated" based on the germination rate, $g$. Survival of the individuals remaining in the seed bank was based on the seed survival rate, $s$. Individuals followed an annual plant life cycle (i.e., individuals who germinated in year $t$ died at the end of year $t$).

#### Simulation process

##### Burn-in

Each simulation began with a burn-in period of 1,000 generations. The goal of this burn-in period was to generate independent replicates of genetically variable focal populations that had evolved in landscapes of similar spatial and temporal heterogeneity for each simulation run. At the start of the burn-in, 10,000 genetically homogenous, perfectly adapted individuals were founded in the central patch of a landscape that contained seven patches. The phenotypic optimum, θ, which determined individual fitness (see *Fitness, mating, and population dynamics* below), changed across the landscape with slope ($b$) = 4.5, with modest, uncorrelated temporal fluctuations [deviations drawn from Normal(0,1)]. This mimics an environmental gradient across the landscape, such as the aridity gradient present across the range of *C. x. xantiana*, and allows for moderate fluctuations in optima through time as would be observed in a natural system. After mating occurred within a population, offspring dispersed according to a Poisson dispersal kernel with mean ($m$) = 0.1. The direction of dispersal (left or right along the gradient) was unbiased and random. This burn-in period allowed mutation, recombination, and gene flow to increase genetic variance of the focal trait, which equilibrated at a high level like those found in our natural populations (mean $V_G$ in focal population at end of burn-in = 12.7; estimated $V_G$ for flowering time in natural populations: KYE = 24.31, S22 = 11.7; Table S4).

##### Main simulation

After the burn-in period, we simulated the 12-year period for which we have long-term demographic and environmental data. Dispersal among populations was halted during this period to avoid any confounding effects of gene flow on evolutionary responses. The central patch in the landscape was considered the focal patch; all summary statistics were calculated using this patch. To simulate the observed demographic patterns in our two populations, carrying capacity during each of the 12 generations of the focal period was proportional to the mean-relativized census size (number of fruiting plants) recorded in those years in our two populations. For the two years at S22 where we recorded zero fruiting plants, we assumed these zeros reflected a non-exhaustive census rather than an actual population size of zero; we set the mean-relativized census size in these years to half of the lowest recorded non-zero value. During the first seven generations of the focal period (corresponding to years 2006-2011 in our observed data), the population phenotypic optimum was equal to the long-term, historical optimum during the burn-in. In the years corresponding to the drought (2012-2015), the phenotypic optimum decreased, with the magnitude of that decrease determined by the model parameterization (see *Simulation model* in main text). Depending on the model scenario (see *Simulation model* in main text), the optimum post-drought (2016-2017) either remained the same or returned to the pre-drought optimum. In each generation, as during the burn-in, the realized phenotypic optimum fluctuated around the set optimum [deviations drawn from Normal(0,1)], in order to include realistic temporal stochasticity in the model.

#### ABC analyses

We used the abc package [7] in R to compare models and estimate parameters. The small number of simulations that went extinct (KYE: 1.3% of simulations; S22: 2.3%) were discarded, as summary statistics could not be calculated. We used the rejection method with a tolerance rate of 0.01 (`method = “rejection”, tol = 0.01`) to calculate posterior probabilities for each model and used the resulting Bayes factors to select the best model for each population. Parameter posterior distributions were estimated using localized linear regression and a tolerance rate of 0.01 (`method = “loclinear”, tol = 0.01`).

### Supplemental tables

**Table S2.** Trait phenotypic means and variances.

| | | KYE | | | S22 | | |
|---|---|---|---|---|---|---|---|
| **Trait** | **Year** | **Mean** | **Variance** | ***n*** | **Mean** | **Variance** | ***n*** |
| *Days to germination* | 2011 | 9.05 | 46.51 | 122 | 10.87 | 36.51 | 83 |
| | 2015 | 9.75 | 33.14 | 104 | 11.16 | 34.55 | 112 |
| | 2017 | 10.52 | 40.68 | 94 | 10.91 | 35.75 | 92 |
| *Growth rate* | 2011 | 0.97 | 0.09 | 94 | 0.95 | 0.11 | 63 |
| | 2015 | 1.08 | 0.11 | 80 | 1.01 | 0.09 | 87 |
| | 2017 | 0.96 | 0.08 | 71 | 1.12 | 0.12 | 70 |
| *Days to flowering* | 2011 | 69.67 | 46.89 | 94 | 69.29 | 28.61 | 66 |
| | 2015 | 69.96 | 26.69 | 80 | 67.67 | 21.25 | 87 |
| | 2017 | 69.84 | 34.69 | 73 | 66.48 | 14.31 | 73 |
| *SLA* | 2011 | 21.28 | 24.43 | 92 | 22.44 | 28.21 | 65 |
| | 2015 | 22.67 | 35.67 | 77 | 22.62 | 28.04 | 84 |
| | 2017 | 20.35 | 16.03 | 71 | 21.12 | 32.80 | 72 |
| *Flowers* | 2011 | 13.84 | 11.74 | 91 | 12.12 | 12.75 | 66 |
| | 2015 | 15.06 | 16.16 | 78 | 12.41 | 16.93 | 88 |
| | 2017 | 13.42 | 15.57 | 72 | 13.48 | 17.50 | 73 |
| *Days to flowering after germination* | 2011 | 60.73 | 31.21 | 94 | 58.12 | 34.08 | 66 |
| | 2015 | 59.81 | 15.24 | 80 | 56.52 | 16.95 | 87 |
| | 2017 | 59.73 | 25.79 | 73 | 55.36 | 16.12 | 73 |
| *Leaf succulence* | 2011 | 5.39 | 0.83 | 92 | 5.42 | 1.05 | 65 |
| | 2015 | 5.52 | 0.84 | 80 | 5.34 | 0.91 | 86 |
| | 2017 | 5.22 | 0.78 | 70 | 5.23 | 1.02 | 72 |
| *Shoot biomass* | 2011 | 501.95 | 17041.45 | 93 | 452.36 | 20052.88 | 66 |
| | 2015 | 491.11 | 20358.95 | 80 | 457.58 | 22034.91 | 88 |
| | 2017 | 470.36 | 17047.90 | 72 | 470.40 | 19369.71 | 72 |

**Table S3.** Results of ANOVA with Type II SS for mixed effect trait models. All models had random effects of sire, and dam nested within sire.

| | | | | | | | | | | | |
|---|---|---|---|---|---|---|---|---|---|---|---|
| | **KYE** | | | | | | | | | | |
| *Fixed effects* | **df** | **Germination** $n$ = 320 | | **Growth rate** $n$ = 245 | | **Days to flowering** $n$ = 247 | | **SLA** $n$ = 240 | | **Flower number** $n$ = 241 | |
| | | $\lambda^2$ | *P* | $\lambda^2$ | *P* | $\lambda^2$ | *P* | $\lambda^2$ | *P* | $\lambda^2$ | *P* |
| Year | 2 | 1.7 | 0.435 | 4.5 | 0.108 | 0.6 | 0.734 | 3.0 | 0.228 | 4.9 | 0.088 |
| Transplant age | 1 | — | — | 1.1 | 0.306 | 91.0 | <0.001 | 0.1 | 0.739 | 10.1 | 0.002 |
| Seed weight | 1 | 18.8 | <0.001 | — | — | — | — | — | — | — | — |
| | **S22** | | | | | | | | | | |
| *Fixed effects* | **df** | **Germination** $n$ = 287 | | **Growth rate** $n$ = 218 | | **Days to flowering** $n$ = 226 | | **SLA** $n$ = 221 | | **Flower number** $n$ = 227 | |
| | | $\lambda^2$ | *P* | $\lambda^2$ | *P* | $\lambda^2$ | *P* | $\lambda^2$ | *P* | $\lambda^2$ | *P* |
| Year | 2 | 0.5 | 0.794 | 3.5 | 0.178 | 10.8 | 0.005 | 4.2 | 0.124 | 2.7 | 0.254 |
| Transplant age | 1 | — | — | 4.6 | 0.033 | 63.1 | <0.001 | 3.5 | 0.063 | 7.6 | 0.006 |
| Seed weight | 1 | 0.5 | 0.461 | — | — | — | — | — | — | — | — |

**Table S4.** Trait additive genetic variance estimates (posterior mean with 95% credible intervals) as estimated by MCMCglmm.

| | $V_A$ | |
|---|---|---|
| **Trait** | **KYE** | **S22** |
| *log(Days to germination)* | **0.15**<br>(0.04 - 0.27) | **0.21**<br>(0.12 - 0.31) |
| *Growth rate* | **0.01**<br>(0.00 - 0.03) | **0.05**<br>(0.03 - 0.08) |
| *Days to flowering* | **24.31**<br>(9.88 - 37.40) | **11.70**<br>(3.96 - 20.27) |
| *log(SLA)* | **0.00**<br>(0.00 - 0.01) | **0.00**<br>(0.00 - 0.01) |
| *Flowers* | **8.97**<br>(2.34 - 18.85) | **11.27**<br>(3.45 - 20.09) |

**Table S5.** Bayes factors (ratios of model posterior probabilities) comparing four models of population demographic and evolutionary dynamics for our two populations. For both populations, Model D was the most probable model.

| | KYE | | | | S22 | | | |
|---|---|---|---|---|---|---|---|---|
| | **A** | **B** | **C** | **D** | **A** | **B** | **C** | **D** |
| *Proportion accepted simulations* | *0.00* | *0.00* | *0.07* | *0.93* | *0.00* | *0.30* | *0.00* | *0.70* |
| **A** (no seed bank; post-drought optimum = pre-drought optimum) | | — | 0.0 | 0.0 | | 0.0 | — | 0.0 |
| **B** (no seed bank; post-drought optimum = drought optimum) | — | | 0.0 | 0.0 | Inf | | Inf | 0.4 |
| **C** (with seed bank; post-drought optimum = pre-drought optimum) | Inf | Inf | | 0.1 | — | 0.0 | | 0.0 |
| **D** (with seed bank; post-drought optimum = drought optimum) | Inf | Inf | 12.9 | | Inf | 2.3 | Inf | |

### Supplemental figures

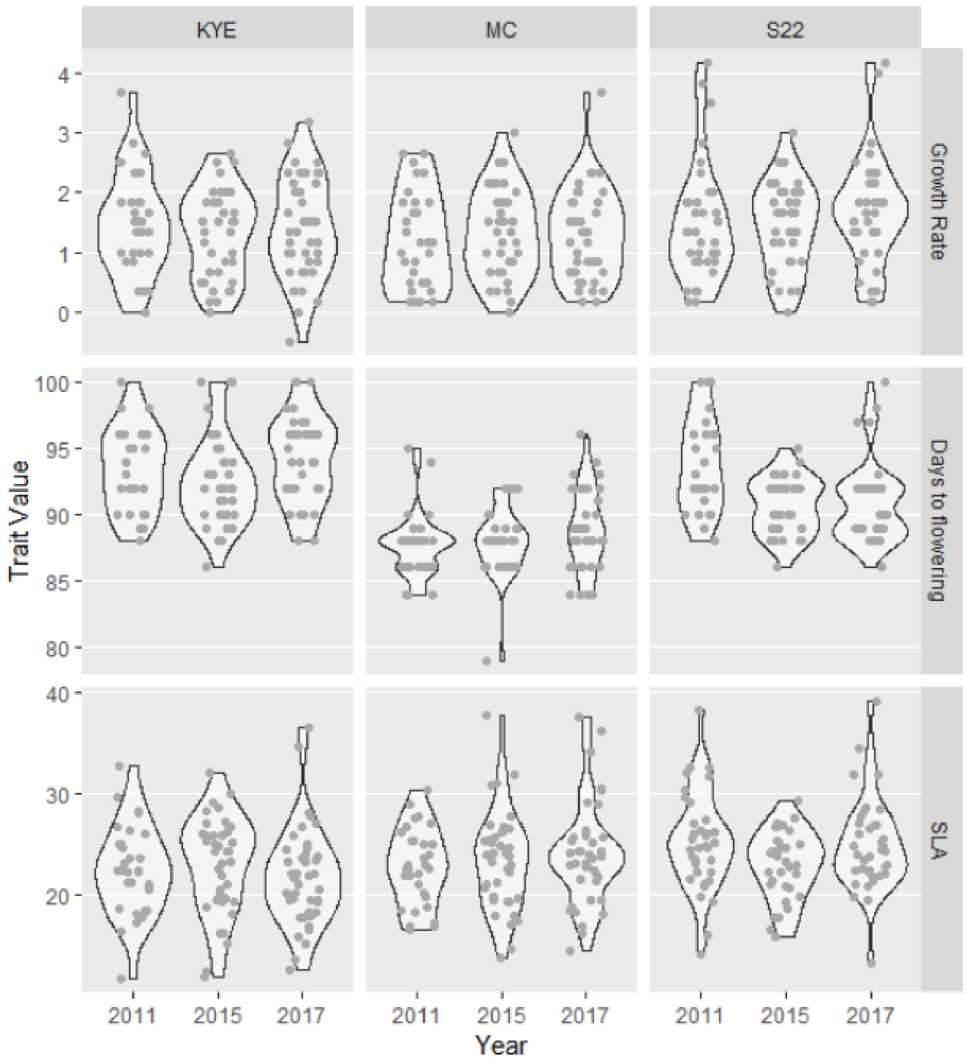


**Figure S4.** Measured phenotypes across the three sampling years for populations KYE, MC, and S22 during the refresher generation.

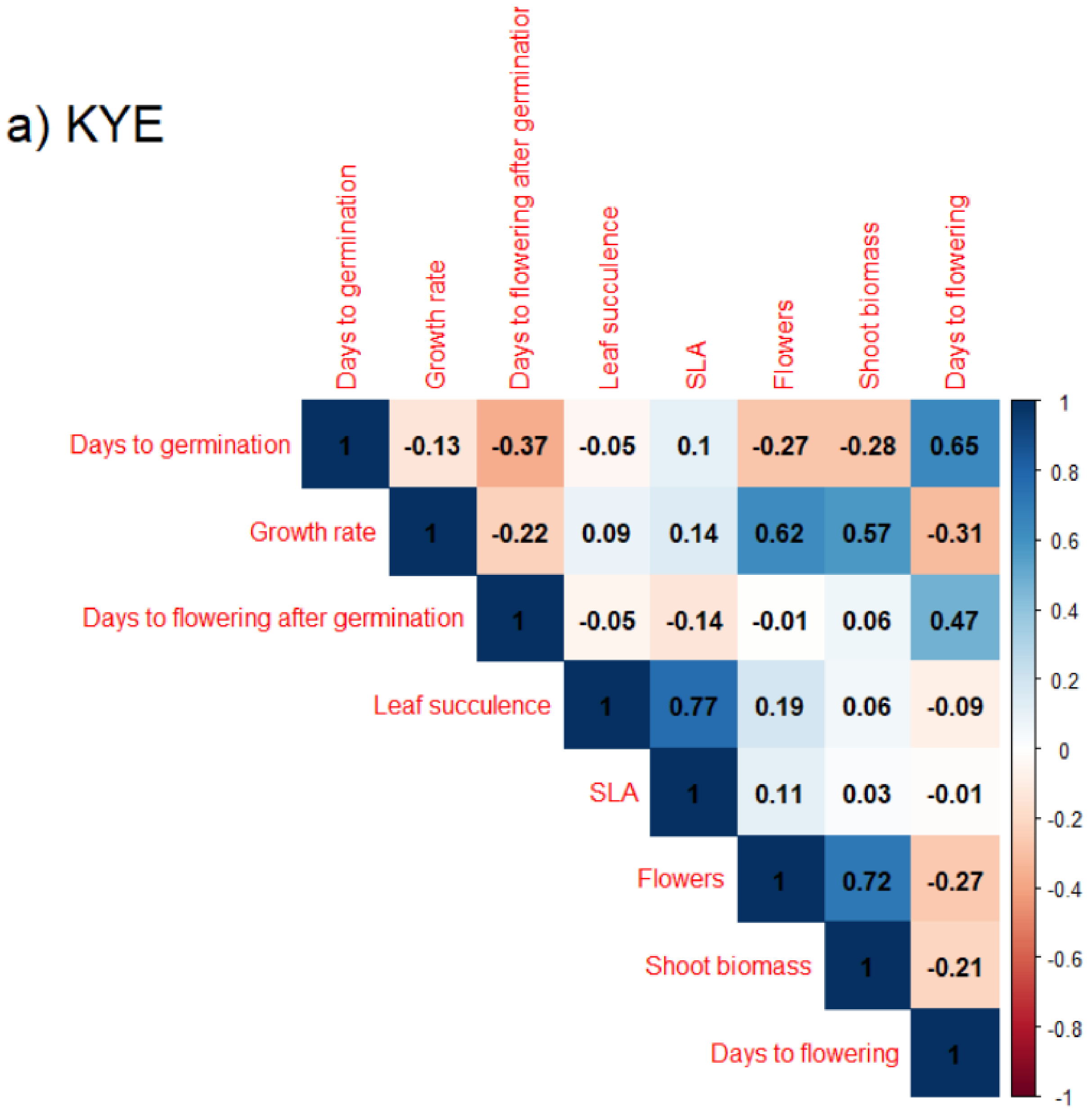

a) KYE
Days to germination
Growth rate
Days to flowering after germinatior
Leaf succulence
SLA
Flowers
Shoot biomass
Days to flowering
Days to flowering after germination
1 -0.13 -0.37 -0.05 0.1 -0.27 -0.28 0.65
1 -0.22 0.09 0.14 0.62 0.57 -0.31
1 -0.05 -0.14 -0.01 0.06 0.47
1 0.77 0.19 0.06 -0.09
1 0.11 0.03 -0.01
1 0.72 -0.27
1 -0.21
1
1
0.8
0.6
0.4
0.2
0
-0.2
-0.4
-0.6
-0.8
-1

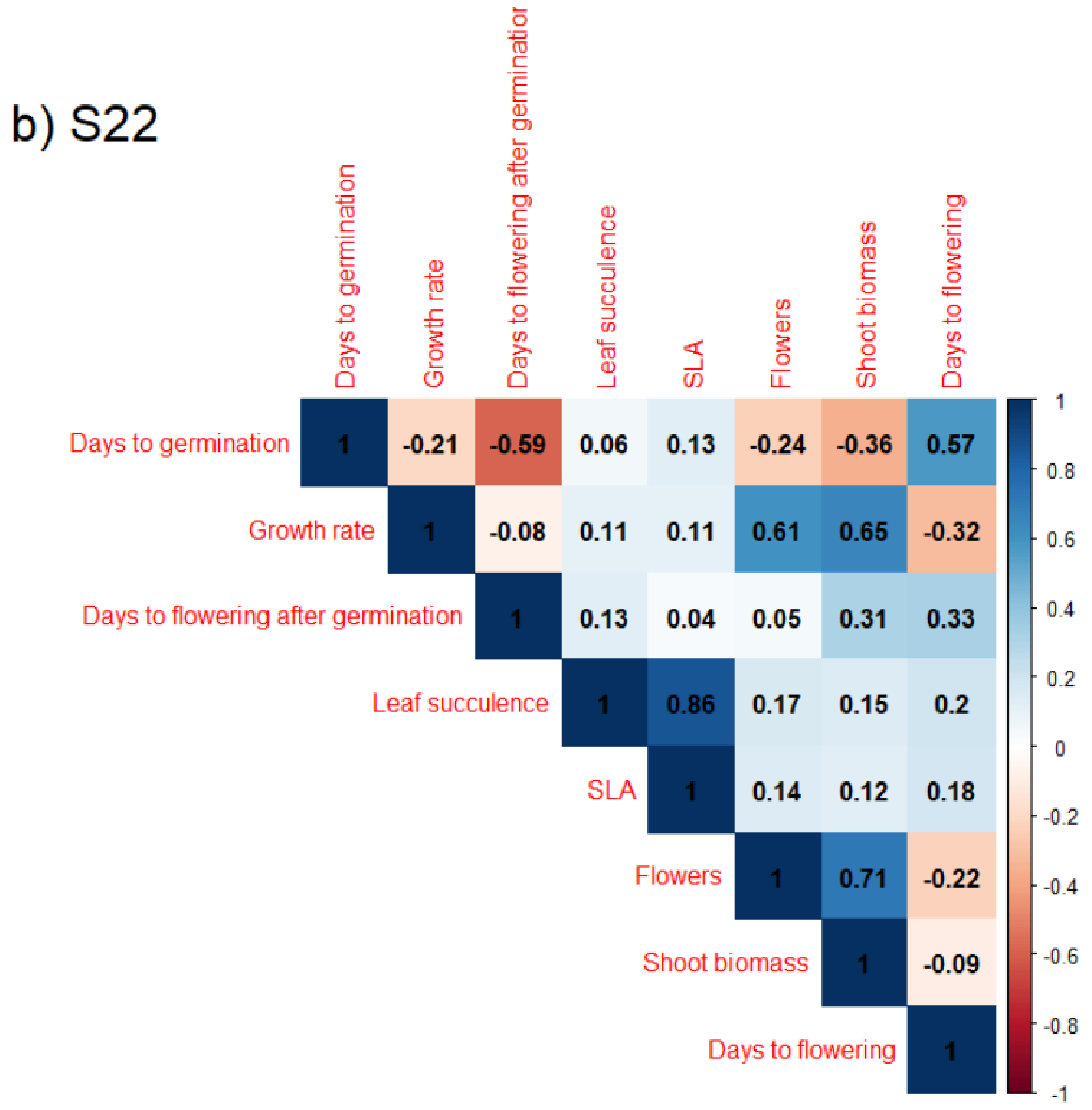


**Figure S5.** Trait correlation matrix for **a)** KYE and **b)** S22.

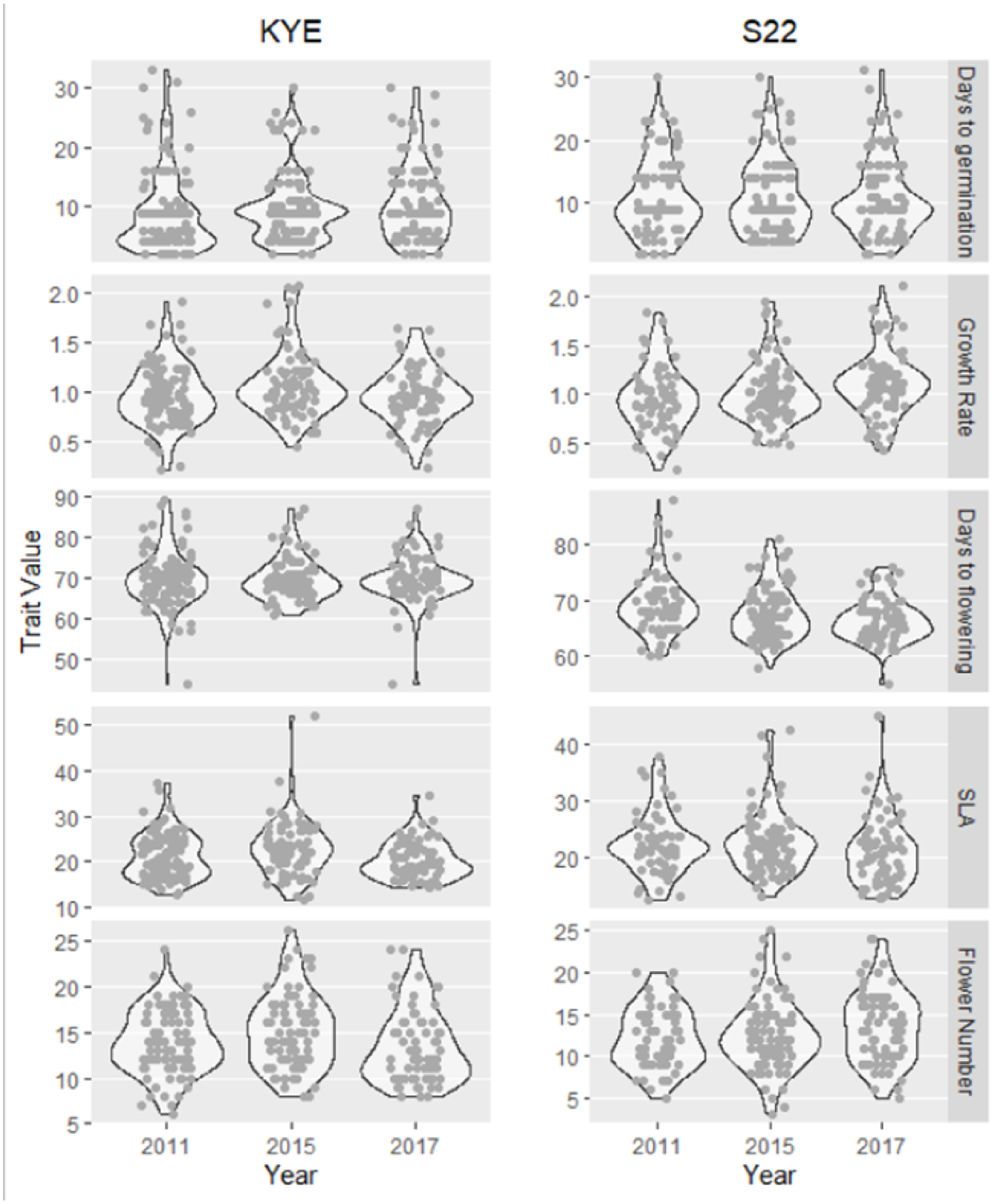


**Figure S6.** Measured phenotypes for both populations in the measurement generation (estimated marginal means shown in Fig. 2).

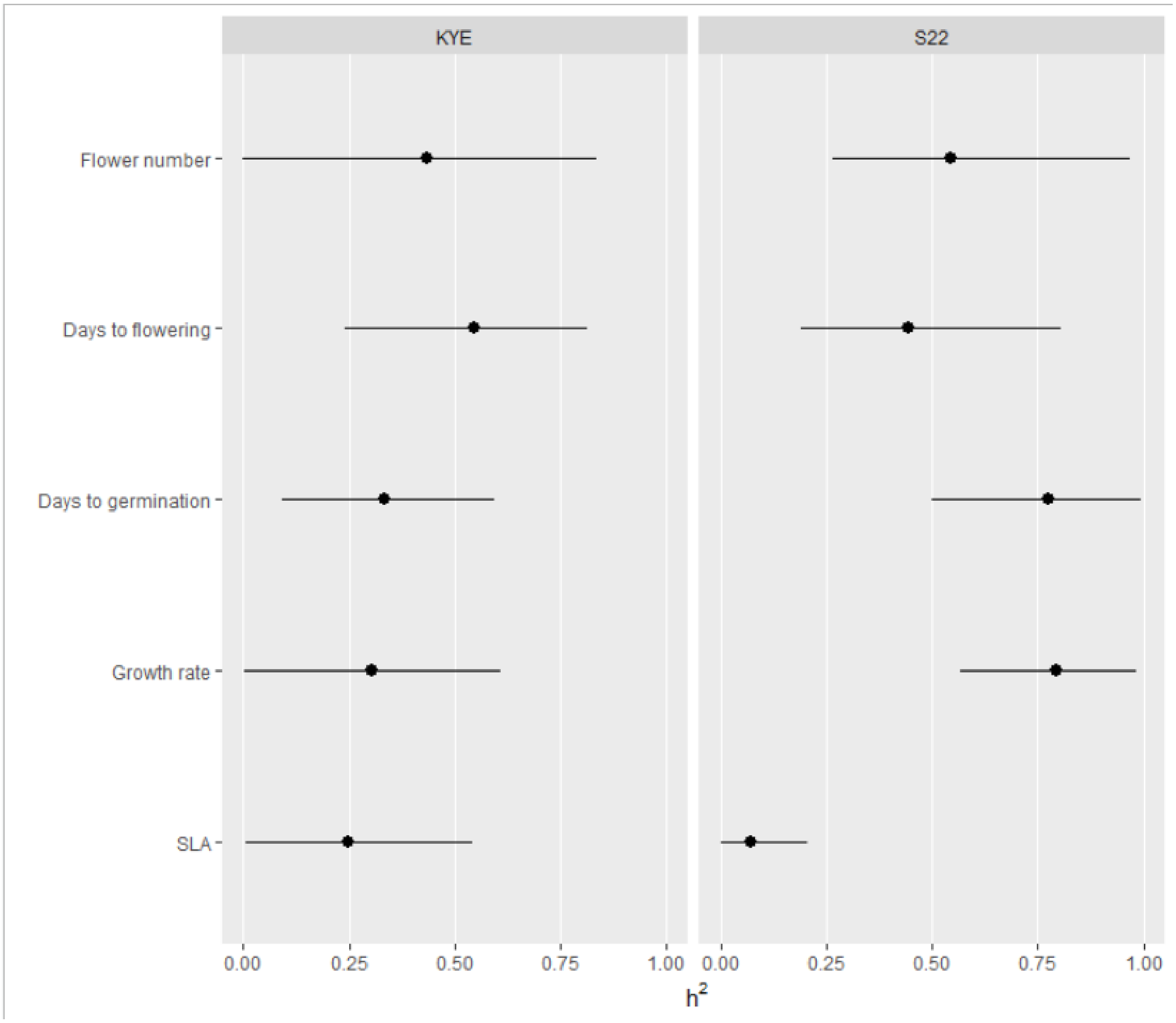


**Figure S7.** Narrow-sense heritability estimates from models using priors without parameter expansion. Compare to Fig. 3 in main text.

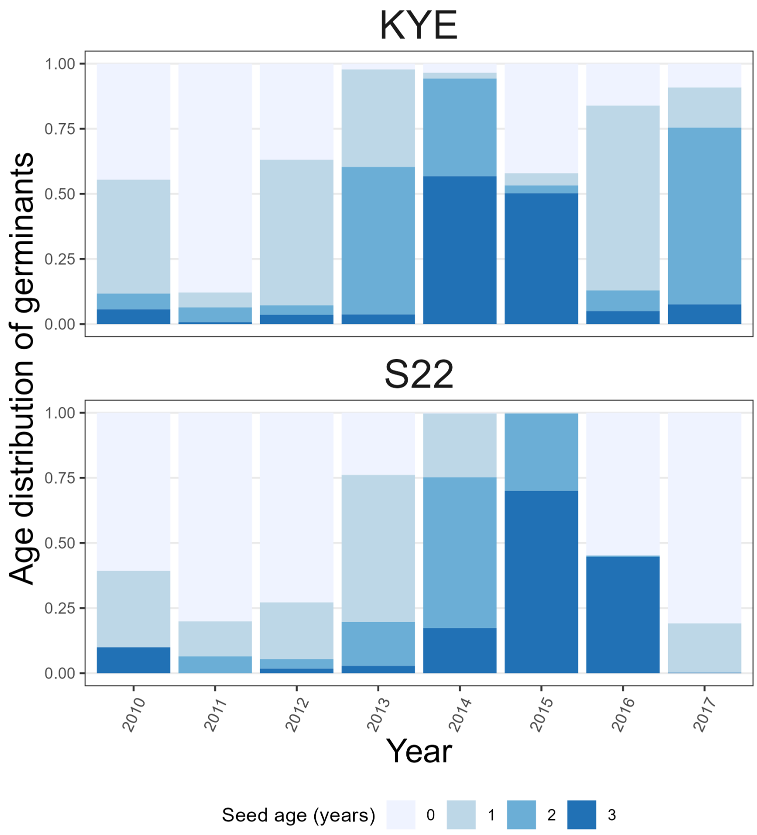


**Figure S8.** Age distribution of germinants in KYE and S22 across years 2010-2017, as estimated based on seed input (present study) and seed bank vital rates from [1].

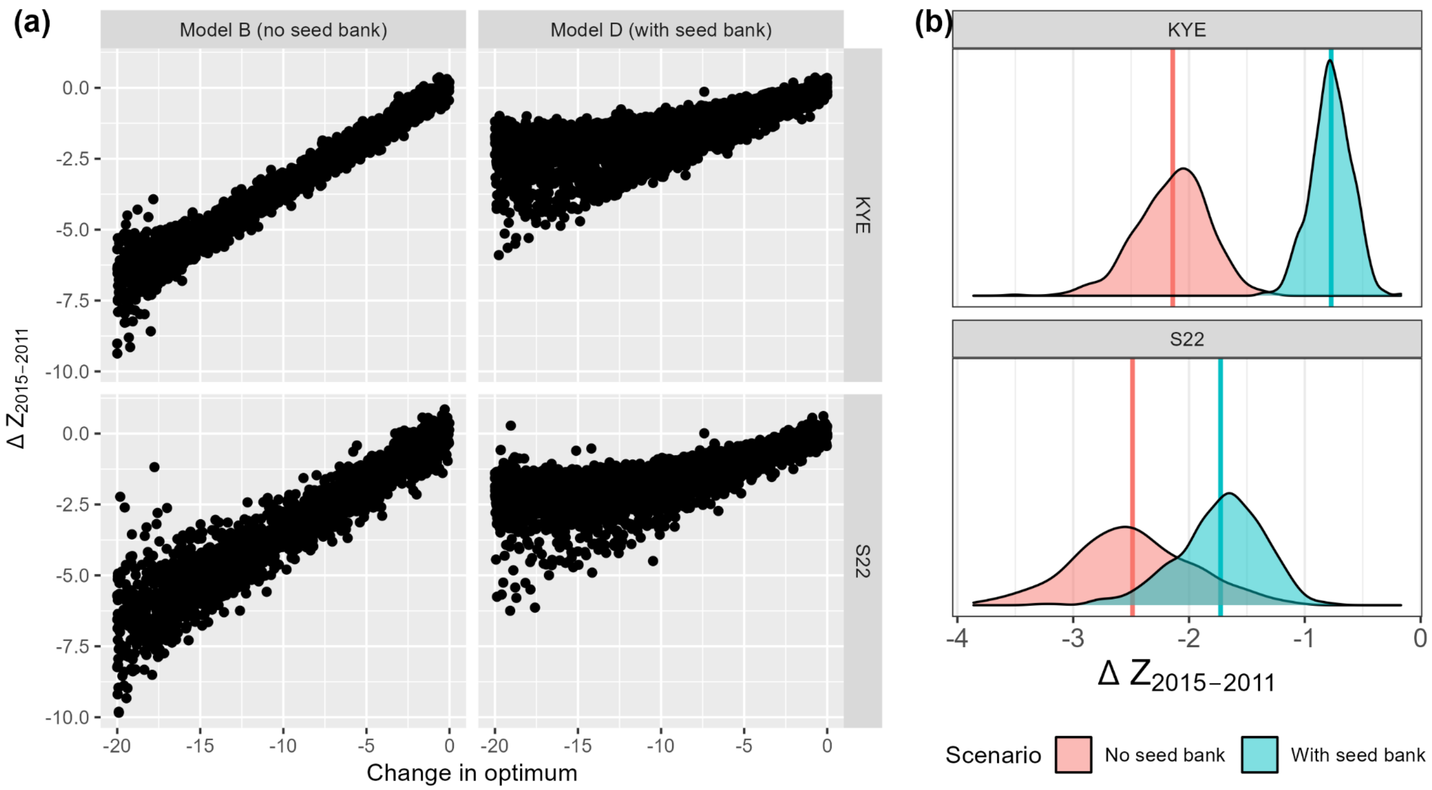


**Figure S9.** Effects of seed banks on phenotypic evolution in our models. (a) Overall effects of the change in phenotypic optimum (x-axis) on the magnitude of phenotypic evolution 2011-2015 in KYE and S22. Each point is a simulation. In the "no seed bank" model (B), germination rate = 1.0 and seed survival rate = 0.0 for all simulations. In the "seed bank" model (D), these vital rates vary from 0.05-0.75 (Table 1). (b) Effect of the seed bank for each population on expected phenotypic evolution 2011-2015. Shown are distributions of the summary statistic $\Delta Z_{2015-2011}$ for 500 simulations run with either the estimated seed bank vital rates for that population ("With seed bank"), or with germination = 1.0 and seed survival = 0.0 ("No seed bank").

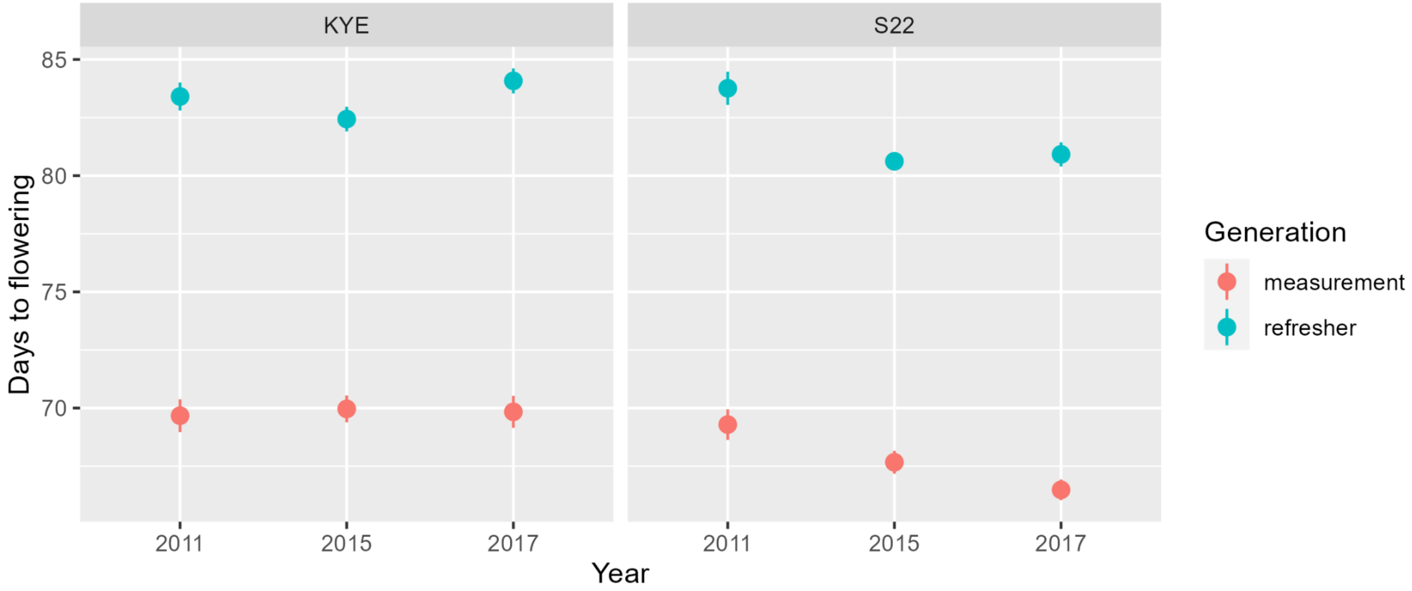


**Figure S10.** The plastic response (vertical distance between blue and red points) in phenology between the refresher (teal) and measurement (red) generations was nearly identical between the two populations. Points show mean ± SE.
